# NAC-mediated ribosome localization regulates cell fate and metabolism in intestinal stem cells

**DOI:** 10.1101/2024.04.29.591601

**Authors:** Sofia Ramalho, Ferhat Alkan, Stefan Prekovic, Katarzyna Jastrzebski, Eric Pintó Barberà, Liesbeth Hoekman, Maarten Altelaar, Rob van der Kammen, Juliette Fedry, Mark C. de Gooijer, William J. Faller, Joana Silva

## Abstract

Intestinal stem cells (ISCs) face the challenge of integrating metabolic demands with unique regenerative functions. Studies have shown an intricate interplay between metabolism and stem cell capacity, however it is still not understood how this process is regulated. Combining ribosome profiling and CRISPR screening in intestinal organoids, we show that RNA translation is at the root of this interplay. We identify the nascent polypeptide-associated complex (NAC) as a key mediator of this process, and show that it regulates ISC metabolism by relocalizing ribosomes to the mitochondria. Upon NAC inhibition, intestinal cells show decreased import of mitochondrial proteins, which are needed for oxidative phosphorylation, and, consequently, enable the cell to maintain a stem cell identity. Furthermore, we show that overexpression of NACα is sufficient to drive mitochondrial respiration and promote ISC identity. Ultimately, our results reveal the pivotal role of ribosome localization in regulating mitochondrial metabolism and ISC function.

**Teaser:** The location of ribosomes in cells is regulated, and defines the fate of intestinal stem cells.

## Introduction

The intestine is a highly dynamic tissue, with a high cell turnover rate. The majority of intestinal cells live for just 4 - 7 days(*1*) before being shed into the intestinal lumen, and intestinal stem cells (ISCs) are central to the maintenance of this epithelium. The fate of ISCs has been shown to be dependent on their metabolic profile, which is typically characterized by high mitochondrial respiration(*2*). Metabolic dysfunction in ISCs results in a loss of stem cells, and can lead to the development of disease(*3*). Furthermore, ISCs are a cell of origin of intestinal cancer(*4*), and considering the crucial role that mitochondrial function plays in cancer maintenance(*5*, *6*), metastasis(*7*, *8*) and acquisition of chemotherapy resistance(*9–11*), a better understanding of the processes that regulate metabolism in ISCs could open the door to future therapeutic interventions. Although several studies have addressed the function of mitochondrial metabolism in these cells(*2*, *12–14*), the molecular mechanism behind this regulation remains largely unknown.

The regulation of mRNA translation plays a critical role in cellular development and function. Although ribosomes have been thought of as passive machines whose function solely consists of passive protein production, recent work from our group and others has shown that this is not necessarily the case(*15*). Studies have demonstrated that ribosomes can exert a direct regulatory function, and that they can play a role as molecular sensors of stress(*16*, *17*). For example, we have shown that in the intestine, ribosomes are crucial in determining the identity of intestinal stem cells (ISCs) in the context of amino acid deprivation(*15*).

In this study, we explore how translation regulates metabolism and consequently stem cell identity, by combining ribosome profiling and CRISPR screening in mouse intestinal organoids. We use two distinct models of metabolic regulation (ISC differentiation and ribosome impairment) to unveil a central role for the nascent polypeptide-associated complex (NAC) in intestinal cell metabolism. NAC is a highly conserved ribosome-associated complex, whose primary function is to interact with newly synthesized polypeptide chains as they emerge from the ribosome during translation, and to assist in their correct folding and targeting(*18*). It is well established that NAC mediates ribosome localization to the endoplasmic reticulum (ER)(*19*), and there have been reports that it can also target ribosomes to the outer membrane of the mitochondria (OMM), however, it is currently unknown whether this OMM targeting is functionally relevant (*20*). Here, we show that it plays a crucial role in defining ISC fate, by facilitating the import of peptides into this organelle, and thus supporting mitochondrial function^20^. Our work goes a step further and establishes NAC as the bridge between translation and metabolism, being translationally regulated itself during ISC differentiation, while also being regulated by ribosome impairment in stress conditions. We further characterize NAC’s role by showing that its effect on metabolism is mediated by the localization of the ribosome to the OMM, and the consequent effect on peptide import. In short, we show that the localization of ribosome plays a central role in ISC maintenance and differentiation.

## Results

### OXPHOS is regulated by mRNA translation in intestinal stem cells

Previous studies have clearly established the importance of metabolic regulation in ISCs. However, some of this work shows that this regulation appears not to be solely at the transcriptional level(*21*). To develop a deeper understanding of the regulators of metabolism in different intestinal cell types, we generated mouse intestinal organoids that were either enriched for stem cells (SCe) or differentiated into enterocytes (SCd)(*22*) (Fig 1A). As expected, the SCe cultures acquired a spheroid morphology and showed an increase in various stem cell markers, such as *Lgr5* and *Axin2*, while differentiated cells show an increase in differentiation markers, such as *Lyz1* and *Alpi* (Fig 1B). We have previously shown that ribosomes play a crucial role in determining the metabolic identity of ISCs in the context of amino acid deprivation(*15*), however the mechanism behind this process, and its importance outside of stress conditions, remains unknown. We therefore assessed the translatome of SCe and SCd organoids using ribosome profiling (RiboSeq), in order to gauge the translational regulation of metabolism in normal intestinal physiology.

**Figure 1.**
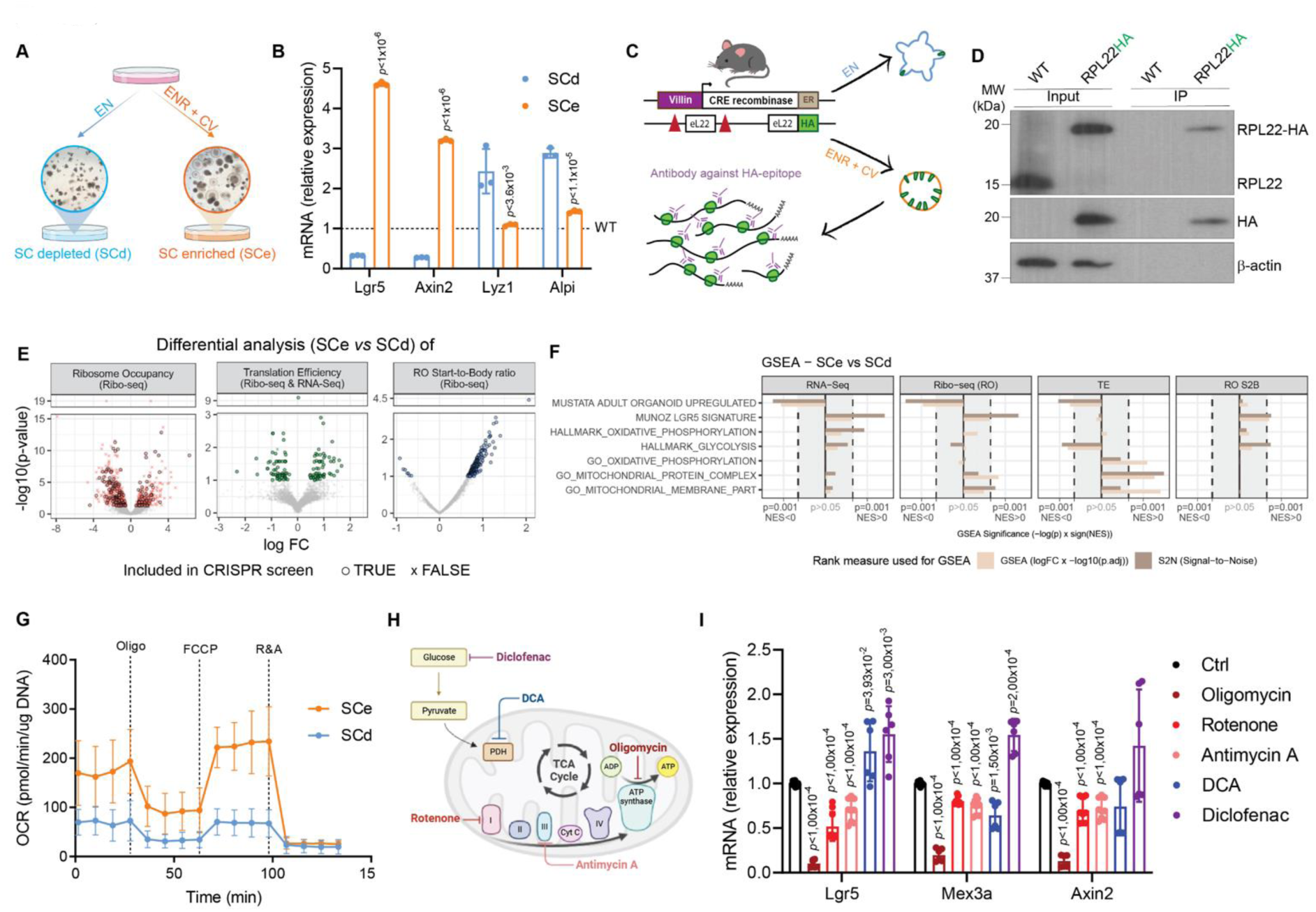
Translation regulation of metabolism affects intestinal stem cell identity. **A)** Schematic figure illustrating the generation of the two in vitro models generated for this study, highlighting the morphological differences between stem cell-depleted (SCd) and enriched (SCe) mouse intestinal organoid cultures. **B)** RT-qPCR analysis of genes related to stem (Lgr5 and Axin2) and differentiation (Lyz1 and Alpi) in SCe and SCd organoids compared to WT cultures (reference dashed line). Hprt was used as a housekeeping control. Mean and SD are shown (n = 3 (technical triplicates for one biological replicate)). p-values were determined using a two-tailed ttest. **C)** Experimental workflow of the RiboSeq experiment developed to compare the translatome of the different cell populations. Intestinal organoids were generated from the VillinCreERT2eL22.HA mouse and grown in SCe or SCd media. HA-tagged ribosomes were pulldown, allowing for the isolation and sequencing of ribosome protected fragments (RPFs). **D)** Western blot analysis confirming efficient pulldown of eL22-HA tagged ribosomes. β-actin was used as loading control and experiments were carried out using one biological replicate. **E)** Volcano plots for the differential analysis of ribosome occupancy (RO), translation efficiency and RO start-to-body ratio (RO S2B) based on RiboSeq and RNAseq data generated with SCe and SCd samples. In all analyses, n = 3 per condition and given p-values in the y-axis are adjusted with the BH method, except for the start-to-body ratio analysis where raw p-values are shown. Genes with significant differences between conditions are highlighted, and genes that are selected for follow-up CRISPR drop-out screening are marked with “+” (Figure 2A). **F)** Combined Gene Set Enrichment Analysis (GSEA) results for a selection of gene sets where separate analyses are performed with standard GSEA and signal-to-noise (S2N) measures. Note that each bar represents an independent GSEA, where genes are ranked based on the given gene expression metric in SCe vs SCd comparison. **G)** Oxygen consumption rate (OCR) analysis shows increased respiration rates in SCe compared to SCd cultures. Mean and SD are shown (n = 5 biological replicates). **H)** Schematic diagram highlighting the metabolic targets of the different OXPHOS and glycolytic inhibitors used in figure 1I. Created with BioRender.com. **I)** RT-qPCR analysis of genes related to stem capacity of WT organoids treated with different OXPHOS (oligomycin, rotenone and antimycin A) and glycolysis (DCA and diclofenac) inhibitors for 24 hours. Hprt was used as a housekeeping control. Mean and SD are shown (n = 6 (two biological replicates, each assessed in technical triplicates)). p-values were determined using a two-tailed t-test.

To do this, we generated SCe and SCd intestinal organoids from mice that express an HA-tagged version of the core ribosomal protein eL22, also known as the RiboTag mouse(*23*, *24*) (Fig 1C). The eL22-HA tag enabled us to efficiently immunoprecipitate ribosomes, from which we could then isolate ribosome protected mRNA fragments (RPFs) for sequencing (Fig 1D, Supp Fig 1). This experiment provided information on ribosomal occupancy both in terms of abundance (ribosome occupancy (RO)) and distribution (RO start-to-body ratio (RO s2b)) for each transcript (detailed in the methods section). Furthermore, by normalizing the RPFs for each transcript with their overall transcript levels, measured by RNA Seq (Supp Figs 2A and B), we were able to identify genes with differential translation efficiency (TE) between SCe and SCd organoids (Fig 1E, Supp Table 1).

**Figure 2.**
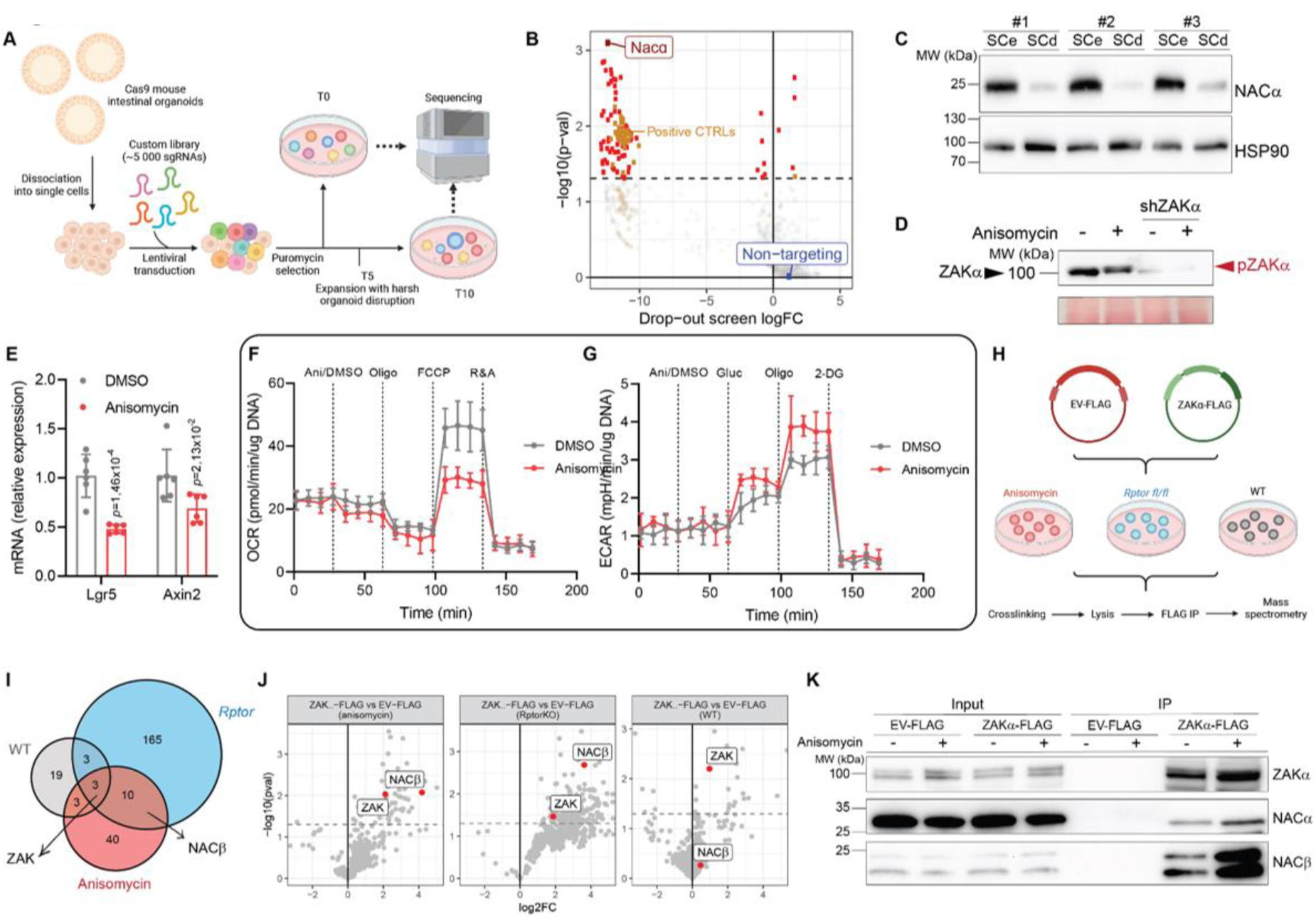
Intestinal Cell Regulation of NAC Occurs Post-Transcriptionally. **A)** Scheme describing the CRISPR dropout screen workflow performed in mouse intestinal organoids and used to assess the essentiality of translationally-regulated hits obtained from previous RiboSeq experiment, for intestinal stem cells (ISCs). Created with BioRender.com. **B)** Results of the CRISPR drop-out screen. Positive controls (genes expected to be important for cell survival), non-targeting gRNA and significant results are highlighted. The top hit is the subunit of NAC, NACɑ. The screen was performed in three biological replicates. **C)** Western blot analysis shows higher levels of NACɑ in SCe compared to SCd organoid cultures. HSP90 was used as a loading control. Experiments were done in three biological replicates. **D)** Immunoblot of Phos-tag gel analysis shows ZAKɑ phosphorylation following anisomycin treatment (1μM, 30 minutes). Experiments were carried out using one biological replicate. **E)** RT-qPCR analysis of genes related to stem capacity of WT organoids treated with 1μM anisomycin for 30 minutes. Hprt was used as a housekeeping control. Mean and SD are shown (n = 6 (two biological replicates, each assessed in technical triplicates)). Pvalues were determined using a two-tailed t-test. **F)** OCR analysis shows decreased respiration rates in WT organoids treated with 1μM anisomycin for 30 minutes. Mean and SD are shown (n = 4 biological replicates). **G)** Extracellular acidification rate (ECAR) analysis shows increased glycolytic rates upon treatment of WT organoids with 1mM anisomycin for 30 minutes. Mean and SD are shown (n = 4 biological replicates). **H)** Schematic figure illustrating the approach used to explore potential new interactors of ZAKɑ in WT mouse intestinal organoids in two different conditions promoting ribosome impairment (Rptorfl/fl organoids and organoids treated with anisomycin), using rapid immunoprecipitation mass spectrometry of endogenous proteins (RIME). An empty vector expressing the FLAG tag bait (EV-FLAG) was used as a control. Created with BioRender.com. **I)** Venn diagram of proteins identified in RIME experiment as potential ZAKɑ interactors in the three different experimental conditions (n = 2 biological replicates). **J)** Volcano plots highlighting differential enriched interactors of ZAKɑ in WT, Rptorfl/fl and anisomycin-treated organoids, using RIME (n = 2 biological replicates). Significance was reached when the p-value < 0,05 (t-test, two-tailed). **K)** Immunoprecipitation of ZAKɑ-FLAG confirms interaction with both NACɑ and NACβ in HCT116 cells, revealing this to be increased upon treatment with 1μM anisomycin for 30 minutes. EV-FLAG-infected cells were used as a control. The experiment was carried out on one biological replicate.

Despite the overall levels of protein synthesis being similar between SCe and SCd cultures (Supp Fig 2C), RiboSeq analysis revealed that around 300 genes are translationally regulated when comparing the two conditions (Fig 1E, Supp Table 1). Most of these hits appear to be regulated by the overall number of ribosomes bound to them (RO). However, we also identified some transcripts that displayed potential changes in initiation and/or elongation rate, as measured by the relative abundance of RPFs at the start of the mRNA compared to the body (RO s2b). Gene set enrichment analysis (GSEA) of this data showed that genesets associated with OXPHOS (and other mitochondrial-associated processes) were changed in ribosome occupancy in SCe organoids compared to SCd, despite no change at the RNA level (Fig 1F and Supp Fig 2B). The opposite trend was observed in gene sets associated with glycolytic pathways, suggesting that this switch may be translationally regulated. Previous studies have demonstrated that ISCs rely heavily on OXPHOS(*2*) and, in accordance with the RiboSeq results, we observed increased levels of mitochondrial respiration in SCe compared to SCd cultures (Fig 1G, Supp Fig 2D). Importantly, when using various OXPHOS inhibitors to impair mitochondrial function, we detected a significant decrease in the expression of stem cell markers (Figs 1H and 1I). In contrast, we saw no consistent effect on the levels of these markers when using different glycolysis inhibitors, supporting the idea that ISCs rely heavily on mitochondrial respiration.

### NACɑ is translationally regulated in ISCs

As we had shown that metabolism is translationally regulated in ISCs, we next assessed the functional importance of translationally-regulated genes in intestinal stem cells. To do this, we performed a dropout CRISPR screen in SCe, Cas9-expressing organoids. We infected these organoids with a custom library containing sgRNAs targeting mRNAs that show altered translational levels in SCe and SCd organoids (Supp Table 2). The library contained a total of 4765 gRNAs, of which 100 gRNAs were non-targeting controls, 445 gRNAs targeted genes known to be important for cell survival (such as ribosomal proteins), acting as positive controls, and 1235 gRNAs were targeting translationally regulated genes involved in metabolic processes. Organoids were cultured in SCe media for 10 days, and midway through this selection (day 5) the organoids were disrupted and recultured, to promote stem cell capacity. Surviving organoids were harvested at day 10 and sequenced, allowing us to identify translationally-regulated genes that are essential for stem cell capacity (Fig 2A). Alongside the positive controls, a total of 74 genes dropped out of the screen. As expected, there was no dropout of the non-targeting guides (Fig 2B, Supp Table 3).

Interestingly, the top hit of the screen was the nascent polypeptide-associated complex subunit alpha, NACα (NACA) (Fig 2B, Supp Table 3). NACα is a subunit of the nascent polypeptide associated complex (NAC), a highly conserved ribosome-associated complex whose primary function is to interact with and assist in the proper folding and targeting of newly synthesized polypeptide chains as they emerge from the ribosome during translation(*18*). We confirmed that NACα levels were changed in SCe compared to SCd organoid cultures by western blotting, which revealed a dramatic post-transcriptional downregulation as ISCs differentiate (Fig 2C, Supp Table 1). This finding suggests a potential role for NACα in ISCs maintenance, potentially via its regulation of metabolic identity.

### ZAKɑ regulates NACβ following translation impairment

We have previously shown that translation impairment also affects the metabolic identity of intestinal cells. This is regulated via the activation of the ribosome stress sensor ZAKα (MAP3K20), driving ISCs from OXPHOS towards glycolysis, which ultimately impacts their identity(*15*). However, how ZAKα mediates this metabolic switch is still unknown. To explore this, we first activated ZAKα in organoids using a low dose of a translation inhibitor (anisomycin) (Fig 2D). This was sufficient to decrease the expression of stem cell markers (Fig 2E) and rewire metabolism by decreasing OXPHOS levels (Fig 2F, Supp Fig 3A) and increasing glycolysis (Fig 2G, Supp Fig 3B). As ZAKα is a kinase, it is likely that it regulates metabolism via direct interaction with a partner. We therefore generated an organoid line expressing a FLAG-tagged version of ZAKα. We treated these organoids with anisomycin, immunoprecipitated ZAKα, and carried out LC-MS (Fig 2H). We compared this to a similar experiment that we previously published(*15*), in which we used the same FLAG-ZAKα to identify interactors following impaired translation via mTOR inhibition. Although both mTOR inhibition and anisomycin treatment activate ZAKα and inhibit OXPHOS, their interactome was surprisingly different (Fig 2I). Nevertheless, analysis of the overlapping interactors identified NACβ (also known as BTF3) as the most significantly enriched common ZAKα-interactor (Figs 2I and J). NACβ is the second subunit of the NAC, forming a hetero-dimer with NACα to assist in the co-translational targeting of nascent polypeptides to the proper organelles.

**Figure 3.**
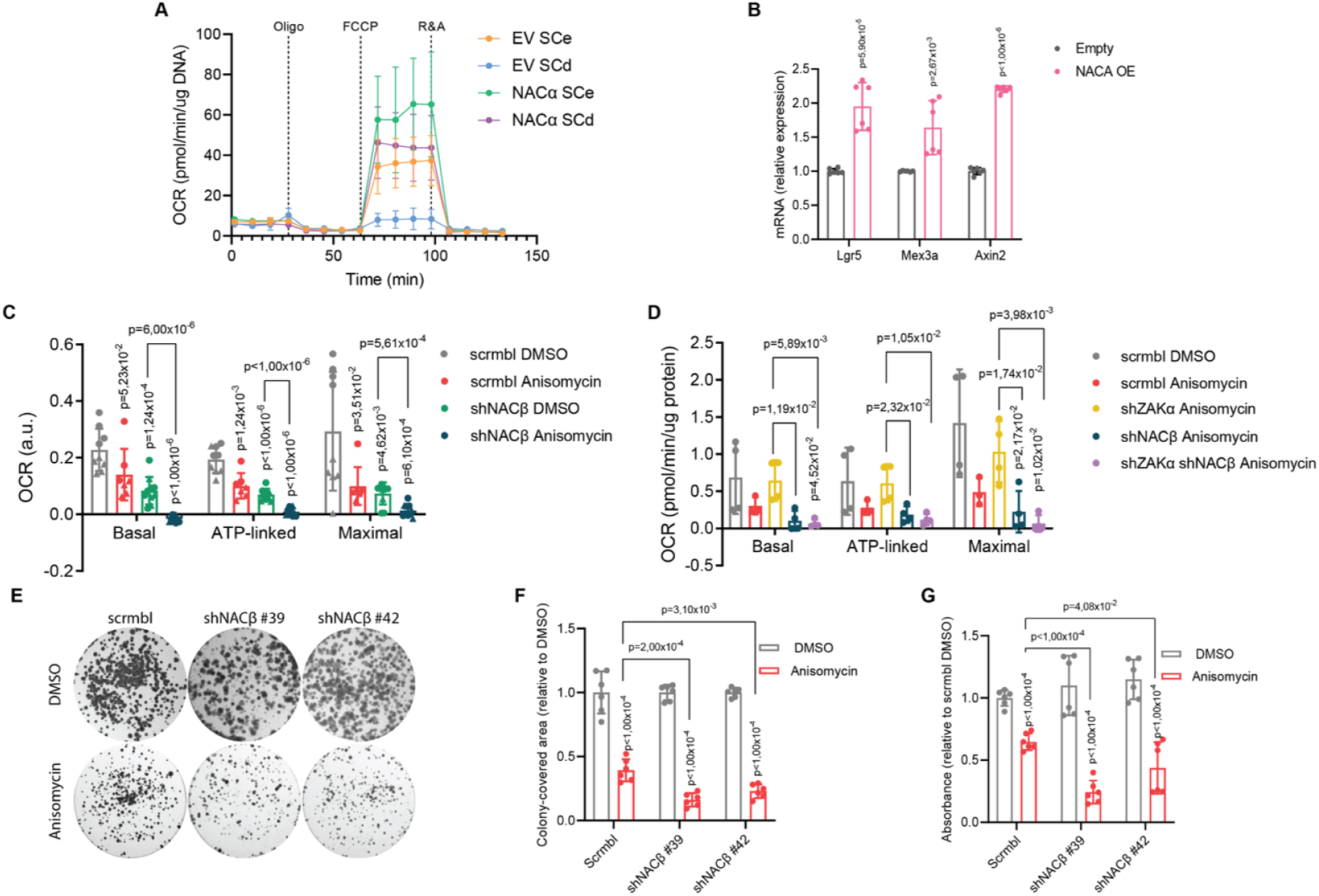
Modulation of NAC affects respiration and stemness in intestinal cells. **A)** OCR analysis shows a rescue of respiration rates in SCd organoids overexpressing NACɑ. Mean and SD are shown (n = 4 biological replicates). p-values were determined using a two-tailed t-test. **B)** RT-qPCR analysis of genes related to stem capacity in organoids overexpressing NACɑ or empty vector control. Hprt was used as a housekeeping control. Mean and SD are shown (n = 6 (two biological replicates, each assessed in technical triplicates)). p-values were determined using a two-tailed t-test. **C)** OCR analysis shows decreased respiration rates in WT HCT116 cells treated with 1μM anisomycin for 30 minutes, upon knockdown of NACβ and when combining both anisomycin treatment with NACβ knockdown. Mean and SD are shown (n = 5 biological replicates for each of the 2 independent shRNAs. Circles mark shNACβ#39 replicates and triangles mark shNACβ#42). p-values were determined using a twotailed t-test. **D)** OCR analysis shows decreased respiration rates in WT HCT116 cells treated with 1μM anisomycin for 30 minutes and the rescue of this decrease when knocking down ZAKɑ. Upon knockdown of NACβ, the rescue of the anisomycin effect observed with the loss of ZAKɑ is not possible. Mean and SD are shown (n = 4 biological replicates). p-values were determined using a two-tailed t-test. **E)** Representative images of survival assay in control (scrmbl) and NACβ knock down HCT116 cells, upon vehicle (DMSO) or anisomycin treatment (1μM, 24 hours). **F)** Quantification of colony-covered area in survival assays shows a decrease in colony covered area upon anisomycin treatment and a further reduction when combined with NACβ knock down. Mean and SD are shown (n = 6 biological replicates). p-values were determined using a two-tailed t-test. **G)** Crystal violet quantification of survival assays, shows a decrease in absorbance upon anisomycin treatment and a further reduction when combined with NACβ knock down. Mean and SD are shown (n = 6 biological replicates). p-values were determined using a two-tailed t-test.

As we had previously identified NACα as a potential regulator of mitochondrial metabolism in ISCs, this obviously piqued our interest. To confirm the interaction of ZAKα with the NAC, we used the colorectal cancer cell line HCT116, in which we overexpressed FLAG-ZAKα. We confirmed that NACβ interacts with ZAKα in these cells, and that this interaction is increased upon anisomycin treatment. Furthermore, we also saw that NACα also interacts with ZAKα, albeit at a lower level than NACβ (Fig 2K).

### Altering NAC levels impacts mitochondrial respiration and intestinal stem cell identity

As we had identified NAC as a potential regulator of OXPHOS in two different model systems, we sought to further explore its role in metabolic regulation in these models and assess whether it plays a role in defining stem cell identity. Using intestinal organoids, we had observed a downregulation of NACα upon cell differentiation (Fig 2C), a process that results in concomitant reduction of OXPHOS (Fig 1G). We therefore hypothesized that overexpression of this gene would promote stemness by upregulating OXPHOS.

This turned out to be the case, and following NACα overexpression in wild-type organoids (Supp Fig 4A), we observed a substantial increase in mitochondrial respiration (Fig 3A and Supp Fig 4B). Moreover, when these organoids were grown in SCd media to promote differentiation, the expected decrease in OXPHOS was completely blocked, emphasizing the importance of NAC in this process (Fig 3A and Supp Fig 4B). Furthermore, NACɑ overexpression also resulted in an increase in stem cell markers (Fig 3B), demonstrating that NAC-mediated regulation of OXPHOS is key in sustaining the metabolic profile necessary for ISC maintenance.

**Figure 4.**
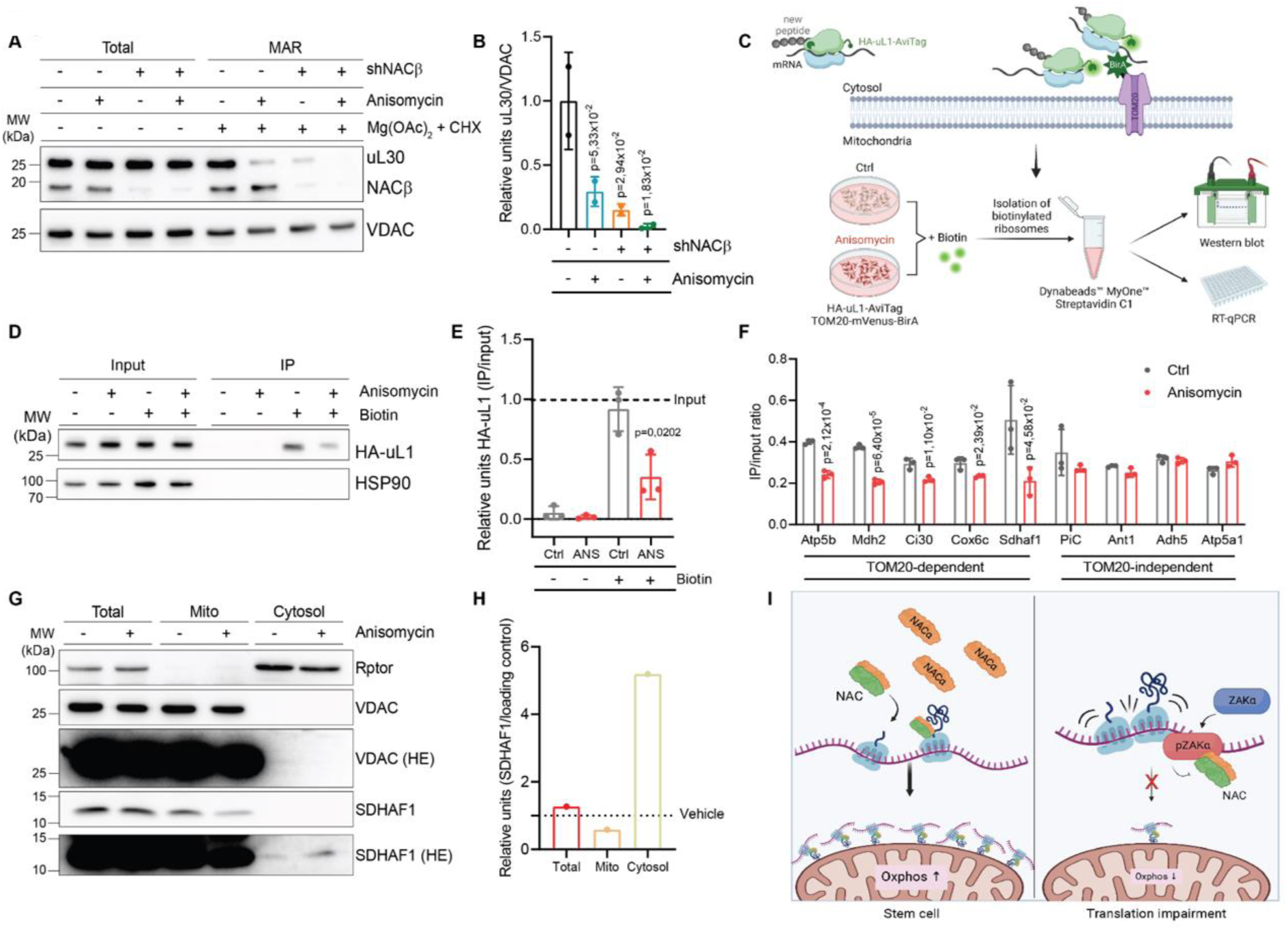
NAC promotes the import of mitochondrial proteins in intestinal cells via ribosome localization. **A)** Western blot showing co-isolation of mitochondrial associated ribosomes (MAR), under stabilization with Mg(OAc)2 and cycloheximide. uL30 blotting shows a decrease of MAR in shNACβ cells and upon treatment with 1μM anisomycin for 30 minutes. VDAC was used as a loading control. Experiments were carried out using two biological replicates. **B)** Western blot quantification (including the one shown in A) shows a decrease of MAR upon anisomycin treatment and knockdown of NACβ. Mean and SD are shown (n = 2 biological replicates). p-values were determined using a two-tailed t-test. **C)** Schematic representation of the ribosome proximty labelling assay used for quantification of ribosomes in proximity of the OMM. HEK293T cells express the OMM protein TOM20 fused to the biotin ligase BirA, which recognizes a specific avidin acceptor fused to the HA-tagged ribosomal protein uL1 (HA-uL1) and biotinilates it upon biotin treatment. Biotinilated ribosomes were isolated using streptavidin beads and further western blot or RT-qPCR analysis was performed. Created with BioRender.com. **D)** Western blot analysis shows a decrease in the number of biotinylated ribosomes, assessed by HA-uL1 levels, following anisomycin treatment. HSP90 was used as a loading control. Experiments were carried out in three biological replicates. **E)** Quantification of western blots (including the one shown in D) shows a decrease in MAR upon anisomycin treatment. Mean and SD are shown (n = 3 biological replicates). p-values were determined using a two-tailed t-test. **F)** RT-qPCR analysis of genes with a putative N-terminal MTS, and therefore imported to the mitochondria in a TOM20-dependent way (Atp5b, Mdh2, Ci30, Cox6c and Sdhaf1) and without, and thus not dependent on TOM20 for mitochondrial import (PiC, Ant1, Adh5 and Atp5a1). 18S rRNA was used as a housekeeping control. Ratios of elute/input, together with mean and SD are shown (n = 3 (technical triplicates for one biological replicate)). p-values were determined using a two-tailed t-test. **G)** Western blot analysis of cell fractionation shows decrease in SDHAF1 levels in mitochondria upon anisomycin treatment. Rptor and VDAC were used as loading controls for the cytosolic and mitochondria fractions, respectively. Experiment was carried using one biological replicate. **H)** Quantification of western blot shown in G shows a decrease in SDHAF1 located in the mitochondria (mito) and an increase in the cytosolic fraction, upon anisomycin treatment. VDAC was used as loading control for total lysate and mitochondria fraction and Rptor for the cytosolic fraction. **I)** Schematic representation of the model proposed for the central role of NAC in metabolic regulation in the intestine. In ISCs there is a translational upregulation of NACɑ which results in an increase in OXPHOS. Upon ribosomal impairment ZAKɑ interacts with NAC, impeding its localization to the OMM and leading to a decrease in OXPHOS. Created with BioRender.com.

Alongside NACɑ overexpression, we also took the opposite approach, and knocked down each subunit of NAC. As this was technically unfeasible in organoids, we used the HCT116 colon cancer cell line model. As expected, based on the CRISPR screen results (Fig 2B), NACα deletion was lethal in these cells. However, we successfully generated NACβ-deficient cells (Supp Figs 4C-E). These cells presented decreased OXPHOS, supporting NACβ’s role as a regulator of metabolism (Fig 3C, Supp Fig 4F). Surprisingly, when combining NACβ deficiency with anisomycin treatment (which results in a ZAKα-mediated decrease in OXPHOS (Fig 3D and Supp Figs 4G and H)), there was an additive effect. We hypothesize that this is a result of incomplete knockdown of NACβ (Supp Figs 4C-E), and that the remaining protein is then fully inhibited by the activation of ZAKα. As expected, anisomycin-mediated inhibition of OXPHOS was rescued by ZAKα knockdown (Fig 3D and Supp Figs 4G and H). Crucially, this rescue was completely abolished in NACβ-deficient cells, suggesting that the effect of ZAKα activation on OXPHOS is mediated by NACβ. Consistent with this, the loss of NACβ also exacerbated the effect of ribosome impairment on the survival of these cells (Figs 3E-G). Using a colony formation assay, we observed that loss of NACβ further decreased both the colony-covered area and the crystal violet staining upon anisomycin treatment, supporting the idea that NAC deregulation decreases OXPHOS and proliferation potential of intestinal cells. Together these results point to the importance of NAC in maintaining the metabolic profile necessary for ISC maintenance, demonstrating that NAC-regulated mitochondrial function plays a crucial regulatory role.

### Mitochondrial protein import is mediated by NAC via ribosome localization

It has been previously shown that NAC can regulate the localization of ribosomes(*19*, *20*). As the local translation of mitochondrial proteins is key to the correct functioning of this organelle(*25*), we reasoned that this may be an explanation of the observed phenotypes. To test this hypothesis, we used a fractionation-based method to measure the abundance of cytosolic ribosomes on the outer membrane of the mitochondria (OMM) (*26*), both in wild-type and NACβ-deficient cells, in the presence or absence of anisomycin. This analysis showed a significant decrease in the number of cytosolic ribosomes associated with the OMM following either anisomycin treatment or NACβ depletion, as measured by uL30 levels (Figs 4A and 4B).

Although this finding supports the idea that impairment of NAC perturbs the localization of ribosomes to the OMM, this experiment relies on the addition of different drugs required to stabilize ribosomes, which may confound the interpretation of these results. To overcome this, we also performed a proximity-labeling assay using cells expressing the OMM protein TOM20 fused to the biotin ligase BirA, which recognizes a specific avidin acceptor fused to the HA-tagged ribosomal protein uL1 (HA-uL1)(*27*) (Fig 4C). After adding biotin to the cells, we immunoprecipitated the biotin-labeled avidin-tagged ribosomes using streptavidin beads and quantified the mitochondrially-associated ribosomes (MAR) by determining the ratio of HA-uL1 levels in IPs-to-Inputs. This confirmed that following anisomycin treatment and consequent ZAKα activation, ribosomes were less localized to the mitochondria (Fig 4D and 4E). As NAC also blocks the mislocalization of proteins to the ER by preventing the signal recognition particle (SRP) from erroneously targeting ribosomes without a signal sequence(*19*), we also measured the distribution of ribosomes to the ER. We employed the same system as above, but using cells expressing HA-uL1-AviTag together with Sec63-BirA(*28*). This showed an increase in ER localized ribosomes following anisomycin treatment (Supp Figs 4I and J). Together, these results indicate that NAC regulates the distribution of ribosomes inside the cell following translation impairment, hampering their targeting to the mitochondria and consequently inhibiting OXPHOS.

To further understand what newly synthesized peptides might be impacted by this MAR relocalization, we isolated the mRNAs associated with biotinylated ribosomes. As it is known that NAC probes the ribosomal exit tunnel for mitochondrial targeting sequence of nascent chains(*18*) which are then imported by TOM20(*29*), we quantified mRNAs that encode for proteins which are thought to be imported by TOM20 (*Atp5b*, *Mdh2*, *Ci30*, *Cox6c* and *Sdhaf1*), compared to mRNAs for mitochondrial proteins that are not thought to depend on TOM20 for their internalization to the mitochondria (*PiC*, *Ant1*, *Adh5* and *Atp5a1*)(*27*). This revealed a significant decrease in the abundance of TOM20-dependent targets upon anisomycin treatment, while transcripts imported by other proteins were unaffected (Fig 4F). Furthermore, through cell fractionation, we analyzed the levels of SDHAF1 in the mitochondria and in the cytosolic fraction. This TOM20-imported peptide is a component of the mitochondrial respiratory chain, and its translation in the vicinity of the mitochondria is reduced upon translation impairment and consequent NAC modulation (Fig 4F). Consistent with this, we observed that upon anisomycin treatment the amount of SDHAF1 in mitochondria decreases, and this is accompanied by an increase in the cytoplasm (Figs 4G and 4H). Together, these results suggest that upon anisomycin treatment, ZAKα interacts with NAC, impairing the localization of ribosomes to the OMM and the consequent import of mitochondrial peptides through TOM20, resulting in the inhibition of mitochondrial metabolism which ultimately affects cell identity in the intestine.

## Discussion

The intestinal epithelium is the primary nutrient sensing organ in the body. It is in direct contact with nutrients and metabolites in the intestinal lumen, and thus needs to constantly adapt to the current conditions by promoting a suitable differentiation program. During these transitions, different intestinal cells display specific metabolic programs, however little is known about how such features are regulated. In this work, we explore how metabolism is regulated in intestinal cells.

It has previously been shown that mitochondrial quality control systems are important for this process, with mitochondrial fission(*14*) and stress response(*13*) playing crucial roles in intestinal stemness. However, our results suggest that this is only part of the picture. We show that ribosomes can modulate mitochondrial function by influencing localized translation at this organelle. Studies have demonstrated a role for localized translation in several contexts; however, these have exclusively focused on the spatial distribution of mRNAs(*30*, *31*). In this study, we show that it is the distribution of the ribosomes that is regulated in ISCs, and that their redistribution is an important part of both normal cell differentiation, and the response to ribosome stress. While it is known that brain cells have the capability to reposition ribosomes during developmental processes and in response to injury(*32*, *33*),our study shows for the first time that ribosome localization itself is a key determinant of cell fate.

We show that this process is mediated by NAC, which is regulated by translation during normal ISC function, and by the kinase Zakα following ribosome stress. In both of contexts, the redistribution of ribosomes acts as a regulator of metabolism and ultimately plays a crucial role in defining their fate. (Fig 4I). By promoting the relocalization of ribosomes to the outside of the mitochondria, NAC ensures that the metabolic needs of ISCs are met. This is a striking observation that brings phenotypic context to the still poorly understood role of NAC-mediated mitochondrial targeting of newly synthetized peptides, and highlights the fact that this complex has a broader role beyond the regulation of SRP recognition of ER signal sequence-containing peptides (*34*, *35*).

This finding also sheds light on the intricate relationship between translation, metabolism and stem cell function. As intestinal cells exhibit a high metabolic plasticity, switching between glycolysis and OXPHOS depending on their specific differentiation needs, the localization of ribosomes to the mitochondrial surface could indicate a specialized translational control mechanism that allows cells to rapidly adapt their protein synthesis in response to different demands. This flexibility is vital for cells to respond to environmental cues and maintain their functions under varying conditions, such as stress-related responses, which happen often in the intestine. Together, we show that the intersection of metabolic cues and signaling pathways determine the fate of ISCs in this complex tissue, and that modulation of RNA translation via ribosome localization is a key mediator of these processes.

It is worth mentioning that while the complexity of the organoid model is a major strength over standard cell line models, it does present some technical constraints. For example, the assessment of ribosome localization was impossible within organoids, necessitating the use of cell lines as an alternative model system for this aspect of our investigation. Nevertheless, the convergence of key phenotypic outcomes between these two distinct experimental approaches lends credibility to our results, providing a compelling rationale for the interpretation of our findings within the broader context of the study’s objectives.

We also show that the metabolic program of ISCs is determined by translation rather than transcription, revealing a new layer of regulation of metabolism in ISCs. This is key to allowing rapid responses that support the metabolic shifts required for self-renewal and/or differentiation into specific cell types. We used RiboSeq to identify this translational regulation, and developed a new metric to analyze this data. This was based not only on ribosomal abundance but also on the distribution of ribosomes along each transcript, a measure we called the RO s2b. This measure provides additional insights into translation dynamics, allowing us to identify mRNAs that have previously been missed by regular RiboSeq analysis. This includes NACɑ, which had a clear difference in RO s2b, but not in overall RO even though its protein expression levels were significantly increased in ISCs. This highlights the power of using different readouts in order to get a more complete understanding of the translational regulation landscape. The combination of this with a custom designed CRISPR screen that was also carried out in intestinal organoids, provided a thorough and reliable analysis of the translational regulation of metabolism.

Our results revealed that ISCs produce high levels of NACα, thus favoring OXPHOS and maintaining their stem cell identity. However, upon either cell differentiation or translational impairment, NAC gets hindered, preventing the targeting of proteins to the vicinity of the mitochondria. This results in a decrease of cellular OXPHOS, and changes in cellular phenotype. Consistent with this, overexpression of NACα is sufficient to rescue the decrease in mitochondrial respiration associated with differentiation, leading to an increase in stem cell markers. Together, these results highlight the complex regulatory roles that translation has in both the metabolism and the identity of intestinal cells. Here, we show for the first time that the intestine is capable of shaping its composition in a quick and adaptable manner by spatially regulating translation of mitochondrial proteins, essential to promote the survival of stem cells. We show that two distinct signals converge on NAC in order to regulate metabolism, positioning it as a central regulator of this process in the intestine. Overall, our findings emphasize the value of considering translation, and particularly ribosome dynamics, in the broader framework of cellular processes, and highlight the importance of the ribosome in understanding the intricate regulatory mechanisms that govern cell identity and function in the intestine.

## Methods

### Ethical approval

All mouse experiments were carried out with the approval of the NKI Animal Welfare Body and the NKI-AVL Institutional Review Board (IRB), according to the ethical and procedural guidelines established by Dutch law.

### Mouse colonies

Experiments in this study included female and male C57BL/6 mice with ages between 8 and 12 weeks old, all bred in-house at The Netherlands Cancer Institute. The animals were generated as described previously(*15*). For *VillinCre^ERT2^*eL22.HA mice, two consecutive injections of 80 mg/kg of tamoxifen were administered, and samples were taken after 24 hours to adjust for variations in recombination efficiency and total cell count. Crypt cultures from *VillinCre^ERT2^Rptor^fl/f^* mice were induced *in vitro*, as described below. *Gt(ROSA)26Sor^tm1.1(CAG-cas9*,-EGFP)Fezh^*mice did not require any induction.

### Organoid isolation and culture

Small intestinal organoids were generated from intestinal crypts of *VillinCre^ERT2^*eL22.HA, *VillinCre^ERT2^Rptor^fl/fl^*and *Gt(ROSA)26Sor^tm1.1(CAG-cas9*,-EGFP)Fezh^* mice, as previously described(*36*). Organoids were cultured in BME (Amsbio) plugs in ENR media: Advanced Dulbecco’s modified Eagle’s media (DMEM)/F12 (Thermo Fisher Scientific), supplemented with 10 mM HEPES (ThermoFisher Scientific), 1x GlutaMAX^TM^ (Thermo Fisher Scientific), 100 U/ml penicillin, 100 µg/ml streptomycin (Thermo Fisher Scientific), 0,1% BSA (Sigma Aldrich), 1x B27 (Thermo Fisher Scientific) and 1x N2 (Thermo Fisher Scientific), together with 10% (v/v) Noggin conditioned media, 10% (v/v) R-spondin conditioned media and 50 ng/ml EGF (Peprotech). All organoid lines were kept at 37°C in a humidified atmosphere with 5% CO2.

For organoids isolated from *VillinCre^ERT2^Rptor^fl/fl^* animals, *Rptor* loss was induced *in vitro*, by treatment with 500nM 4-hydroxytamoxifen (Sigma Aldrich).

Stem cell-enriched cultures (SCe) were generated by culturing organoids in ENR media supplemented with 10 µM CHIR99021 (Cayman) and 1,5 mM of valproic acid (VPA, Biovision)(*22*) for 4-5 days. Stem cell depleted cultures (SCd) were obtained when culturing organoids in EN media (same media described above, but depleted of R-spondin) for 2 days.

In order to inhibit oxidative phosphorylation, the organoids were treated for 24 hours with 2,5 µM oligomycin A, 1 µM rotenone or 1 µM antimycin A. For glycolysis inhibition, organoids were treated for 24 hours with 0.5 mM diclofenac or 15 mM dichloroacetate. All drugs used in the Seahorse analysis were purchased from Sigma Aldrich.

### Cell culture

HCT116 and HEK293T were obtained from the American Type Culture Collection (ATCC). HEK293T HA-uL1-AviTag and HA-uL1-AviTag Sec63-mVenus-BirA cells were a kind gift from Jonathan S. Weissman lab (UCSF)(*28*) and Yoav Arava lab (Technion)(*27*). All cells were cultured according to standard methods in Dulbecco’s modified Eagle’s media (DMEM) High Glucose, Glutamax-supplemented media (Thermo Fisher Scientific) supplemented with 10% fetal calf serum (FCS), 100 U/ml penicillin and 100 µg/ml streptomycin (Thermo Fisher Scientific), at 37°C in a humidified atmosphere with 5% CO2. All cell lines used were routinely tested for mycoplasma contamination.

### Plasmids, lentiviral production and infection

LentiGuide-Puro (Addgene)(*37*) vector was used for cloning the custom CRISPR library used in the screen. shRNAs constructs for knockdown of NACβ were chosen from the Open Biosystems Expresion Arrest^TM^ TRC library (target sequences: sh#39 CCCAGCATCTTAAACCAGCTT and sh#42 GCAGCGAACACTTTCACCATT). The target sequences were inserted into a pLKO.1 vector (Addgene). As a negative control an shRNA containing a scramble sequence (ACAAGATGAAGAGCACCA) expressed in the same vector backbone was used. For the knockdown of ZAKɑ the commercial construct pLV[shRNA]-Puro-U6>hMAP3K20[shRNA#5] (VB221006-1160bjs, VectorBuilder) was used (target sequence GATGTGACATTCAACACTAAC). The scramble shRNA lentiviral control vector pLV[shRNA]-EGFP/Puro-U6>Scramble_shRNA (VB010000-0009mxc, VectorBuilder) was used as a control. For proximity assays, plasmid containing Tom20-mVenus-BirA was a kind gift from the Yoav Arava lab (Technion)(*27*). This construct was transiently expressed in cells using polyethylenimine (PEI, Polysciences) as a transfection agent (3 µl/DNA µg).

Lentivirus was produced by transfecting the above-mentioned vectors into HEK293T cells with the third-generation lentiviral packaging plasmids pVSV-G, pRSV-REV and pMDL RRE (Addgene). PEI (Polysciences) was the transfection agent used for plasmid delivery (3 µl/DNA µg). The viral supernatant was filtered through a 0,45 µm filter and, in the case of organoid infection, was concentrated using the LentiX Concentrator reagent (Takara).

For the affinity purification and immunoprecipitation experiments, ZAKɑ-FLAG was expressed through the construct pLV[Exp]-Bsd-mPGK>mMap3k20[NM_023057.5]/3xFLAG (VB210329-1381j, VectorBuilder) and the empty FLAG vector LV[Exp]-Bsd-mPGK> {3xFLAG/Stuffer_300bp} (VB210329-1386ht, VectorBuilder) was used as a negative control. For NACɑ overexpression the commercial construct pLV[Exp]-Bsd-CMV>mNaca[NM_013608.3]/FLAG (VB230413-1266frg, VectorBuilder) was used. These constructs were packaged into lentivirus by VectorBuilder. Detailed information can be retrieved on vectorbuilder.com using the identifiers.

Cells were infected with lentivirus using 8 µg/m polybrene (Sigma Aldrich) and were subsequently selected with 2 µg/ml puromycin (Thermo Fisher Scientific) or 8 µg/ml blasticidin (Thermo Fisher Scientific) depending on the infected vector.

In organoid infections, organoids were kept in enriched ENR media: ENR media supplemented with growth and stem-cell inducing factors 10 µM Rho kinase inhibitor Y-27632 (Cayman), 1 mM VPA (Biovision), 1 µM Jagged-1 (AnaSpec) and 6 µM CHIR99021 (Cayman), for 2 days. Organoids were then dissociated into single cells using 1x TryplE Express Enzyme solution (Thermo Fisher Scientific) or StemPro Accutase (Thermo Fisher Scientific), in the case of *Rptor fl/fl* organoids. Cells were plated over BME-coated wells in enriched ENR media and infected with the concentrated virus using 8 µg/m polybrene. 24 hours after infection, a layer of BME was put on top of these cells and the media was refreshed. Then the organoids were selected with either 2 µg/ml puromycin (Thermo Fisher Scientific) or 8 µg/ml blasticidin (Thermo Fisher Scientific) depending on the infected vector. Modified organoids were then cultured in BME plugs with ENR media.

### RNA isolation and quantitative real-time PCR (RT-qPCR)

Cells were collected on ice, washed in PBS and centrifuged at 1000rpm for 5 minutes at 4°C. Pellets were lysed with TRIzol^TM^ Reagent (Thermo Fisher Scientific). RNA was isolated by chloroform extraction followed by centrifugation, isopropanol precipitation, washing in 75% ethanol and resuspension in nuclease-free water. Nucleic acid quantification was performed with Nanodrop and 1 µg of template was used for downstream analysis. Reverse transcription reactions were carried out using the High-Capacity cDNA Reverse Transcription Kit (Thermo Fisher Scientific), following manufacturer’s instructions. qPCR was performed using the comparative CT method by normalization of targets of interest to a suitable housekeeping gene (Table 1). SYBR^TM^ Green PCR Master Mix (Thermo Fisher Scientific) reactions were carried out in technical triplicates in a final volume of 7 µl.

### Protein synthesis assay

Protein synthesis rates were measured as described previously(*36*). Briefly, intestinal organoids were grown in either SCe or SCd conditions and taken on day 4 for analysis. Cells were incubated with DMEM methionine-free media (Thermo Fisher Scientific) for 20 minutes, after which 30 µCi/ml ^35^S-methionine label (Hartmann Analytic) was added for 1 hour. Cells were harvested in ice-cold PBS and centrifuged at 800rpm for 3 minutes to clear the BME. Pellets were resuspended in lysis buffer (50 mM TrisHCl pH 7.5, 150 mM NaCl, 1% Tween-20, 0.5% NP-40, 1x protease inhibitor cocktail (Roche) and phosphatase inhibitor cocktail (Sigma Aldrich) and incubated on ice for 10 minutes. Lysates were then cleared by centrifugation at 13000 rpm for 2 minutes and precipitated onto filter paper (Whatmann) with 25% trichloroacetic acid and washed twice with 70% cold ethanol and twice with cold acetone. Finally, a liquid scintillation counter (Perkin Elmer) was used to measure scintillation and the activity was normalized by total protein content. All experiments were done in technical replicates for each biological unit.

### Bioenergetics analysis

Assessment of the cellular metabolic function was carried out using Seahorse Bioscience XFe24 Analyzer (Agilent). Measurements in organoids were done as described previously by our group(*15*). For HCT116 cells, these were seeded at 25 000 cells/well and cultured in DMEM, as described above, for 48 hours prior to the analysis. Both oxygen consumption rate (OCR) (representing mitochondrial respiration) and extracellular acidification rate (ECAR) (representing glycolysis) measurements were taken according to the manufacturer’s instructions in DMEM (Sigma Aldrich) supplemented with 2 mM L-glutamine for the ECAR experiments and additional 5,5 mM D-glucose for the OCR measurements. For OCR assays, the following reagents were added: oligomycin A (1 µM), FCCP (1 µM) (Sigma Aldrich), rotenone (1 µM) and antimycin A (1 µM). For the ECAR analysis the following reagents were added: glucose (10 mM) (Sigma Aldrich), oligomycin A (1 µM) (Sigma Aldrich) and 2-deoxy-D-glucose (2-DG) (50 mM) (Sigma Aldrich). When stated, cells were pre-treated with either anisomycin (1 µM, Sigma Aldrich) or vehicle (DMSO) for 30 minutes. Results were normalized to DNA content for the organoids. Briefly, cells were dissolved in DNA lysis buffer (75 mM NaCl, 50 mM EDTA, 0.02% SDS, 0.4 mg/ml Proteinase K) and incubated at 56°C for 2 hours. DNA was then precipitated by mixing samples with 1 volume of isopropanol, followed by an incubation at 4°C overnight, and centrifugation at 8000 rpm for 30 minutes at 4°C. Pellets were washed with cold 70% ethanol and air dried. Finally, DNA was resuspended in nuclease-free H2O and quantified using Nanodrop. For cell lines, total protein content was used to normalize the results. Cells were lysed in RIPA lysis buffer (50 mM Tris-HCl pH 8.0, 150 mM NaCl, 1% NP-40, 0,5% sodium deoxycholate, 0,1% SDS, 1x protease inhibitor cocktail cOmplete^TM^ ULTRA Tablets, EDTA-free (Roche)), incubated for 10 minutes on ice and centrifuged for 20 minutes, max speed at 4°C. Supernatants were quantified using the Pierce^TM^ BCA Protein

Assay Kit (Thermo Fisher) and an Infinite^Ⓡ^ M Plex microplate reader (Tecan).

### Western Blot

Organoids and cells were washed twice with cold PBS and pellets were resuspended on ice in the appropriate lysis buffer. Samples were then sonicated for 10 cycles of 1 second ON/1 second OFF, with an amplitude between 20% to 30%. After quantifying proteins with the Pierce^TM^ BCA Protein Assay Kit (Thermo Fisher) and an Infinite^Ⓡ^ M Plex microplate reader (Tecan), these were separated by SDS-PAGE and transferred to a 0.2 µm pore nitrocellulose membrane (PALL). Membranes were blocked using 5% milk/TBS-T (Tris-buffered saline 0,1% Tween 20), then incubated with primary antibodies overnight at 4°C followed by the corresponding secondary antibody conjugated to horseradish peroxidase (HRP), for 1 hour at room temperature. Finally, proteins were visualized with the help of ECL-Plus reagent (Thermo Fisher) and Syngene equipment. The antibodies used in this study can be found in Table 2.

### Phos-tag gel electrophoresis

Cells were lysed in 2x NuPAGE^TM^ LDS sample buffer (Thermo Fisher) with 100 µM DTT, sonicated and boiled for 10 minutes at 100°C. Samples were run in 8% SDS-PAGE gels prepared with 10,7 µM Phos-tag^TM^ acrylamide (Wako) and 21,4 µM MnCl2. Before transfering, the gel was rinsed three times with transfer buffer supplemented with 10 mM EDTA and then one final time with EDTA-free transfer buffer. Transfer and subsequent procedure was performed as described for western blot.

### RNA Sequencing

Total RNA was isolated from SCe and SCd organoid cultures as described above. The quality of the samples was assessed with the 2100 Bioanalyzer, using an RNA Nanochip (Agilent), and used in downstream analysis when showing an RNA integrity number (RIN) above 8. Libraries were generated with the TrueSeq Stranded mRNA kit (Illumina) and sequenced using HiSeq2500 equipment.

### Ribosome profiling

#### Sample preparation

Samples were prepared as described previously(*15*). Briefly, intestinal organoids were generated from *VillinCre^ERT2^*RPL22.HA and *VillinCre^ERT2^Rptor^fl/fl^*RPL22.HA mice and plugged in 30 μl of BME. Each experimental condition was carried out using 3 biological replicates and around 150 plugs were used for each replicate. Translating ribosomes were stalled by treating cells with 100 μg/ml cycloheximide for 3–5 minutes at 37 °C and immediately incubating them on ice for the remainder of the experiment. After collecting the cells, pellets were washed twice with cold PBS supplemented with 100 µg/mL cycloheximide, resuspended in ice-cold lysis buffer (20 mM Tris HCl pH 7.4, 10 mM MgCl2, 150 mM KCl, 1% NP-40, 100 µg/mL cycloheximide and 1x EDTA-free proteinase inhibitor cocktail (Roche)) and incubated for 20 minutes on ice, followed by mechanical disruption with a 25G syringe. Lysates were then centrifuged at max speed for 20 minutes at 4 °C and the supernatants were collected.

#### eL22-HA pull down

Lysates were precleared with Pierce™ Control Agarose Matrix (Thermo Fisher Scientific) for 20 minutes at 4°C, and immunoprecipitated with pre-washed AntiHA.11 Epitope Tag Affinity Matrix (BioLegend) for 4 hours at 4°C. Beads were washed twice with ice-cold lysis buffer and twice with ice-cold wash buffer (20 mM Tris HCl pH 7.4, 10 mM MgCl2, 350 mM KCl, 1% NP-40, 100 µg/mL cycloheximide and 1x EDTA-free protease inhibitor cocktail (Roche)). Tagged ribosomes were then eluted by incubating the beads with 200 µg/mL HA peptide (Thermo Fisher Scientific) for 15 minutes at 30°C with constant agitation. Digestion of non-protected RNA was performed with 10 µl of RNase I (Thermo Fisher Scientific) for 40 minutes at 25 °C and the reaction was stopped by adding 13 µl of SUPERASE (Thermo Fisher Scientific). Lastly, ribosome-protected fragments (RPFs) were purified using miRNeasy minikit (Qiagen), following manufacturer’s instructions.

#### Library preparation

The library preparation was performed as previously described(*15*). RPFs ranking from 19 to 32 nucleotides were size-selected using a 10% TBE-Urea polyacrylamide gel. 3’ and 5’ ends were modified accordingly and the respective adapters were ligated with a T4 RNA ligase I (New England Biolabs). The final products were size-selected one last time and rRNA depletion was performed using custom-made biotinylated oligos(*38*) (Table 3) together with MyOne Streptavidin C1 DynaBeads (Thermo Fisher Scientific). Purified RPFs were then used to synthesize cDNA with SuperScript III (Thermo Fisher Scientific), according to manufacturer’s instructions, using the RTP primer (Table 1). After purification with G50 columns (Merck), cDNA was amplified using Phusion High-Fidelity DNA Polymerase (Thermo Fisher Scientific), with primers containing different RP indexes to allow for sequencing (Table 1). PCR products were purified with a QIAquick PCR purification kit (Qiagen) and size-selected one last time with an E-Gel SizeSelect II 2%, (Thermo Fisher Scientific). The quality and molarity of all samples were assessed with the Agilent 2100 and libraries were sequenced on the Illumina HiSeq2500.

#### RNA Seq and RiboSeq Data analysis

All sequencing datasets were first quality controlled using the FastQC tool. Ribosome profiling data was later trimmed and cleaned from adapter sequences using the Cutadapt tool(*39*). RiboSeq-related QC plots were generated with RiboCode(*40*) after mapping the rRNA&tRNA-cleaned reads to mm10 genome using the STAR aligner(*41*). Transcript quantifications were performed with Salmon (v.1.1.0)(*42*), for which transcript sequences from gencode vM21 annotation were used. However, CDS and UTR sequences of each gene were separated for the quantification of Ribosome Occupancy (RO) with RiboSeq reads. Prior to transcript quantifications, sequencing reads were also cleaned from rRNA fragments with the SortMeRNA tool(*43*). In subsequent analyses, uORF RO was defined as the ribosome occupancy levels determined on the 5’UTR sequences of each gene. Differential translation efficiency of each gene was calculated with the RiboDiff tool(*44*), for which the input consisted of salmon-computed RNA Seq-based mRNA abundance and RiboSeq based CDS Ribosome Occupancy values for every primary transcript of each gene. Primary transcripts were determined by the APPRIS annotation and low-abundance genes were excluded from this analysis. The other gene-specific translation metric, RO start-to-body (S2B) ratio, was defined as the ratio of read counts between the translation initiation site (first 30 codons) and the rest of the CDS, where each gene is represented by its primary transcript. Note that this ratio is also multiplied by the CDSlength-90 to minimize the effect of varying CDS lengths on RPF abundance, and genes/transcripts with a low number of RPFs or CDS length smaller than 100 are excluded from the analysis. Differential analysis of RNA abundance, RO and uORF RO were performed with the DESEQ2 package(*45*) in the R environment, whereas, *limma* package(*46*) was used for differential S2B ratio analysis. Gene Set Enrichment Analyses (GSEA) were performed with the ClusterProfiler package(*47*) using two different ranking measures: the standard GSEA measure and signal-to-noise ratio (S2N). The former measure was defined as logFC x min(-log10(adj p-value),3), except for the differential S2B ratio where standard *p*-values were used instead of adjusted ones. S2N was defined as the difference of group-mean values between conditions divided by the sum of standard deviation in both groups(*48*).

### CRISPR screen

#### Library design

To assess the essentiality of translationally regulated genes in intestinal stem cells, we created a customized CRISPR library where target genes/regions were selected based on analysis specific criteria and manual selection of control targets. From our analyses, we included all genes with significant TE changes (adj.P<0.1) or S2B ratio changes (p.val<0.1) to be targeted in the CDS region. Similarly, genes with significant RO change (adj.P<0.05) but with no significant change in expression (adj.P>0.05) were also targeted at the CDS region. Genes with significant uORF RO change (adj.P<0.05) were targeted in three separate regions; CDS, uORF (UTR5) and upstream region from the uORF. For all CDS regions, we selected 5 guides from Brie(*49*) and GeCKO(*50*) libraries, whereas, for each UTR5 and UP regions, we designed 5 highly efficient and specific guides with the help of CRISPRon(*50*) and CRISPRoff(*51*) web servers.

#### Custom library cloning

Custom library was ordered from Twist Biosciences. Esp3I recognition sites were appended to each sgRNA sequence together with appropriate overhangs sequences and adaptor identifiers to allow differential amplification from the ordered synthesis pool by PCR with primers *PC1* (Table 1). The PCR product was purified using the MinElute PCR Purification kit (Qiagen). Amplicons were cloned into the LentiGuide-Puro vector (digested according to the lentiviral CRISPR toolbox protocol developed by Zhang lab(*37*) (Addgene’s website) and purified by agarose gel electrophoresis using the Wizard^Ⓡ^SV Gel and PCR Clean-Up System (Promega) via Golden Gate cloning with FastDigest Esp3I (Thermo Fisher Scientific) and T7 DNA ligase (3000U/µl) (New England Biolabs). The product was purified and transformed into Endura^TM^ DUOs Electrocompetent cells (LGC, Biosearch Technologies) according to manufacturer’s instructions. DNA was purified using the PureLink^TM^ HiPure Plasmid Filter Maxiprep kit (Invitrogen) and equal sgRNA coverage within the library was confirmed by sequencing.

#### Library transduction and selection

Lentivirus was produced and Cas9 expressing organoids were transduced at 600-fold complexity of the library at a multiplicity of infection (MOI) of 0.3 for each of the three technical replicates. The precise working virus titre was determined in advance in a viral titration experiment on non-Cas9 expressing organoids in which survival was measured following puromycin selection. 60×10^6^ cells were plated on a BME layer, infected with the determined viral volume and 24 hours after transduction their growth into organoids was promoted by overlaying them with BME. A day after this puromycin selection started (2 µg/ml) and after 48 hours selection of infected organoids was completed and T0 triplicates were collected. The replicates of the experimental arm were cultured for 10 days in ENR media supplemented with 10µM CHIR99021 (Cayman) and 1,5mM of VPA (Biovision). On day 5 organoids were harshly disrupted by pipetting in order to potentiate their stem capacity. The surviving organoids were collected on day 10.

#### Barcode amplification and sequencing

DNA of all time points and replicates was extracted using the DNeasy Blood and Tissue kit (Qiagen). sgRNA cassettes were amplified from the collected genomic DNA and Illumina sequencing adaptors and sample barcodes were introduced in two consecutive PCR reactions, first with primers *PCR1* and then with *PCR2* primers (Table 1), using Phusion polymerase (2U/µl) (Thermo Fisher Scientific). PCR products were purified with a QIAquick PCR Purification kit (Qiagen), molarity was assessed with the Agilent 2100 and samples were sequenced on the Illumina NextSeq550.

#### Data analysis

Adapter trimming of sequencing data was performed with cutadapt(*39*). Quality control and drop-out analysis was performed with MAGECK-VISPR pipeline(*52*).

### Affinity purification and quantitative mass spectrometry

Rapid immunoprecipitation mass spectrometry of endogenous proteins (RIME) was used to identify potential ZAKɑ interactors as previously described(*15*, *53*). In addition to the wild-type and *Rptor^fl/fl^* organoids used in our previous work(*15*), we also included one condition of wild-type organoids treated with 1µM of anisomycin for 30 minutes. The exact same conditions were used to immunoprecipitate ZAKɑ from HCT116 cells, with western blot analysis being the main readout to test interaction partners.

### Survival Assay

HCT116 cells were plated in 6 well plate at around 1500 cells/well and cultured in DMEM as described above, for 10 days. Upon plating cells were either treated with anisomycin (1 µM, Sigma Aldrich) or vehicle (DMSO) for 24 hours. After 10 days, cells were fixed and stained with crystal violet solution (0,025% crystal violet (Sigma Aldrich), 1% MetOH, 1% formaldehyde) for 1 hour and then washed with water. Images were acquired in a ChemiDoc XRS+ (BioRad) and colony-covered area was quantified using Fiji software. Crystal violet staining was extracted by incubating plates with 10% acetic acid solution for 40 minutes and 562 nm absorbance was measured using an Infinite^Ⓡ^ M Plex microplate reader (Tecan).

### Mitochondria-associated ribosome (MAR) isolation

MAR samples were isolated as previously described(*26*) with a few alterations. Briefly, cells were treated with 50 µg/ml cycloheximide and 2 mM Mg(OAc)2, upon either anisomycin (1 µM) or vehicle (DMSO) treatment for 30 minutes. Cells were then harvested in extraction buffer (0,25 M sucrose, 20 mM HEPES-KOH pH=7.5, 10 mM KCl, 1,5mM MgCl2, 1 mM EDTA, 1 mM EGTA, 1 mM DTT, 0,1 mM PMSF, 50 µg/ml cycloheximide and 2 mM Mg(OAc)2) and homogenized on ice using a Teflon glass apparatus (loose pestle) 20 times. After centrifugation at 750 g for 10 minutes at 4°C, supernatants were collected and the pellets were resuspended in extraction buffer and once again homogenized and centrifuged with the same settings. Both supernatants were combined and centrifuged at 10000 g for 15 minutes at 4°C. Pellets containing crude mitochondria were resuspended in sucrose/MOPS buffer (250 mM sucrose; 10 mM MOPS-KOH pH=7.2, 50 µg/ml cycloheximide and 2 mM Mg(OAc)2) and proteins were solubilized in laemmli buffer with 50 mM DTT, at 65°C for 15 minutes, and analyzed by western blot. A list of the antibodies used can be found in Table 2.

### Proximity assays

HEK 293T cells expressing HA-uL1-AviTag, Tom20-mVenus-BirA and/or Sec63-mVenus-BirA were grown in DMEM media (Thermo Fisher Scientific) supplemented with 10% (v/v) dialyzed fetal calf serum (dFCS) and 5 0U/ml penicillin/50 µg/ml streptomycin. Cells were treated with 100 µg/ml of cycloheximide (Sigma Aldrich) for 2 minutes at 37°C, followed by labelling with D-biotin (50 µM) (Sigma Aldrich) for 20 minutes at 37°C. After washing with PBS supplemented with 100 µg/ml cycloheximide, cells were collected by scraping in lysis buffer (20 mM Tris pH 7.5, 150 mM NaCl, 5 mM MgCl2, 2% Triton X-100, 1 mM DTT, 100 µg/ml cycloheximide) and lysed on ice for 5 minutes. Lysates were then cleared by centrifugation at 3000 g for 15 minutes at 4°C and the supernatants were loaded on a Zeba de-salt spin column (Thermo Fisher Scientific). Proteins were quantified using the Pierce™ BCA Protein Assay Kit (Thermo Fisher Scientific) and 100 µl were saved for input control. 2 mg of total protein extracts were incubated with 50 µl pre-washed Dynabeads™ MyOne™ Streptavidin C1 beads (Thermo Fisher Scientific) for 1 hour at 4°C using an overhead tumbler. The supernatant was removed with the help of a magnetic rack and the beads were washed three times with a high salt wash buffer (20 mM Tris pH 8.0, 750 mM KCl, 5 mM MgCl2, 0.1% Triton X-100, 0,5 mM DTT, 100 µg/ml cycloheximide) for 15 minutes at 4°C. Biotinylated proteins were eluted from the beads with 10 U of TEV protease (Invitrogen), according to the manufacturer’s instructions. For western blot analysis, 10% of input and 25 µl of Strep-IP elute were analyzed by Western Blot as described above. For RT-qPCR analysis, the protocol was the same as described, but the elution was done by adding Trizol to the beads and proceeding with the RNA isolation, cDNA synthesis and real-time q-PCR, as described above, using the primers listed in the Table 1.

### Cell fractionation

Cells were lysed in STM buffer (250 mM sucrose, 50 mM Tris-HCl pH7.4, 5 mM MgCl2) on ice for 30 minutes. After centrifugation at 4°C, 800 g for 15 minutes, supernatants were collected and and centrifuged again, for 10 minutes. Sepernatants were once more centrifuged at 4°C for 10 minutes, this time at 11 000 g, the pellets and supernatants were collected for mitochondria and cytosolic isolation, respectively. Pellets were resuspended in 200 µl STM buffer and centrifuged at 11 000 g for 10 minutes, 4°C. The resulting pellets were resuspended in 100 µl SOL buffer (50 mM Tris-HCl pH 6.8, 1 mM EDTA, 0,5% Triton X-100) and sonicated three times for 10 seconds, avoiding warm up. For the cytosolic fractions, the supernatants were precipitated in acetone for 1 hour at -20°C and centrifuged at 4°C, 12 000 g for 5 minutes. Pellets were resuspended in 300 µl STM buffer.

### Data availability

The sequencing data for SCe organoids has been previously published(*15*) and can be accessed through the NCBI Gene Expression Omnibus repository, using the <u>GSE180208</u> accession id. Sequencing data for SCd organoids and CRISPR screen has been deposited in the same repository, and available with the id <u>GSE230307</u>. The proteomic datasets have been already included in a previous publication(*15*) and uploaded to the PRIDE repository with the <u>PXD033122</u> accession id. Source data are provided with this paper.

## Acknowledgements

We thank Dr. Kevin B. Myant, Dr. Patrizia Cammareri, Dr. Adam E. Hall (IGC, Scotland), Dr. Roderick Beijersbergen and Dr. Ben Morris (NKI, The Netherlands) for insightful discussions regarding the customized CRISPR screen in intestinal organoids; Dr. Fabricio Loayza-Puch (DKFZ, Germany) for helping develop the RiboSeq protocol for intestinal organoids; and Dr. Jonathan S. Weissman (UCSF, United States) and Dr. Yoav Arava (Technion, Israel) for the cells and plasmids used in the proximity assay. Work in the Faller lab is supported by the KWF (NKI-2021-13878), and the NWO (OCENW.KLEIN.263). JS is supported by an EMBO Long Term Fellowship [210-2018]. JF is supported by an NWO Veni fellowship (212.152).

## Contributions

S.R., W.J.F. and J.S. conceptualized the study, designed experiments and analyzed the data. F.A. performed all the bioinformatic analyses. S.R., S.P., E.P.B., R.vd.K. and J.S. carried out experiments. S.P., K.J., J.F. and M.C.d.G. helped design experiments and provided intellectual input to the manuscript. L.H. and M.A. performed mass spectrometry experiments. S.R., W.J.F. and J.S. wrote the manuscript with input from the other authors. W.J.F. and J.S. supervised experiments.

---

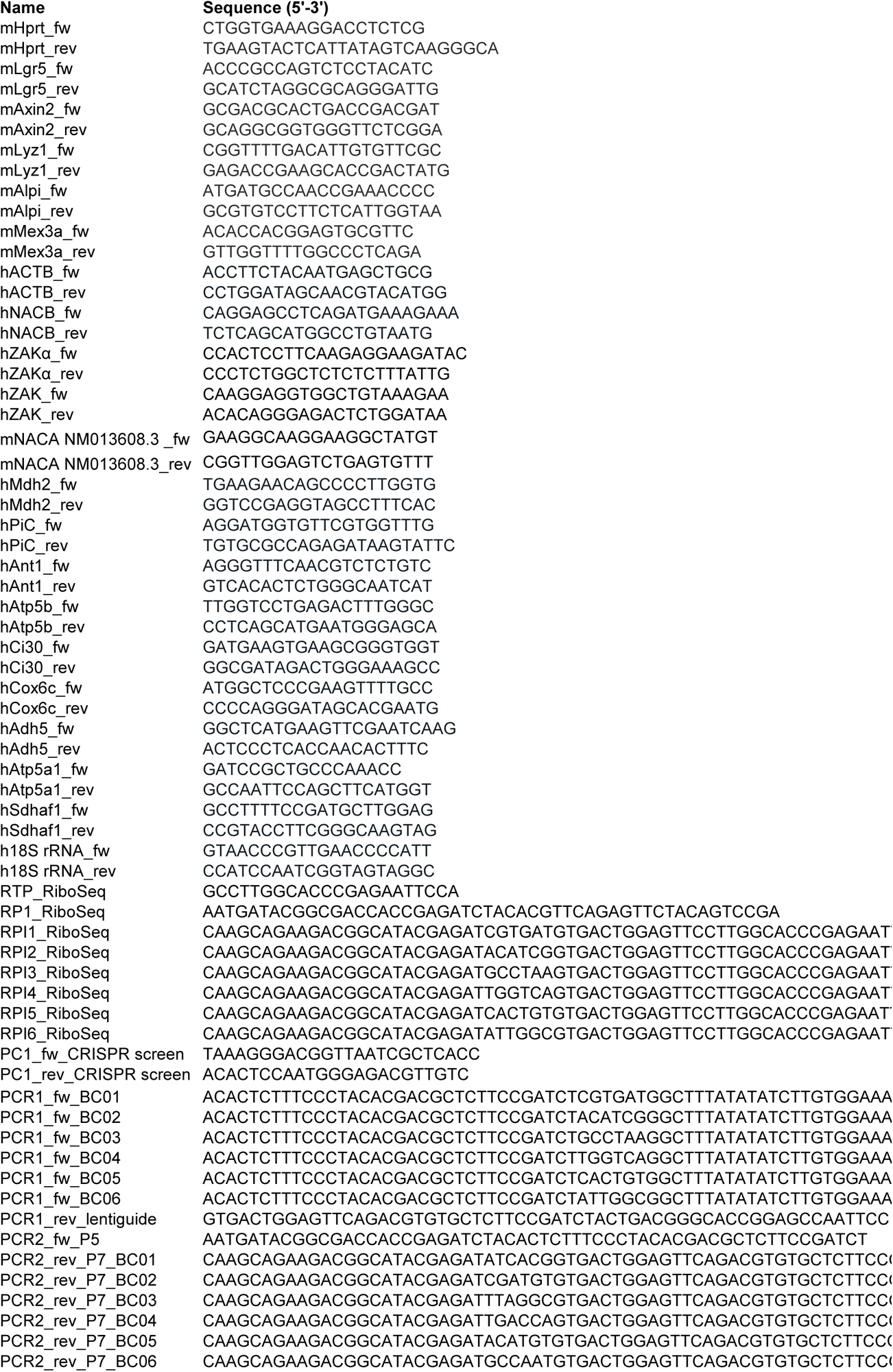

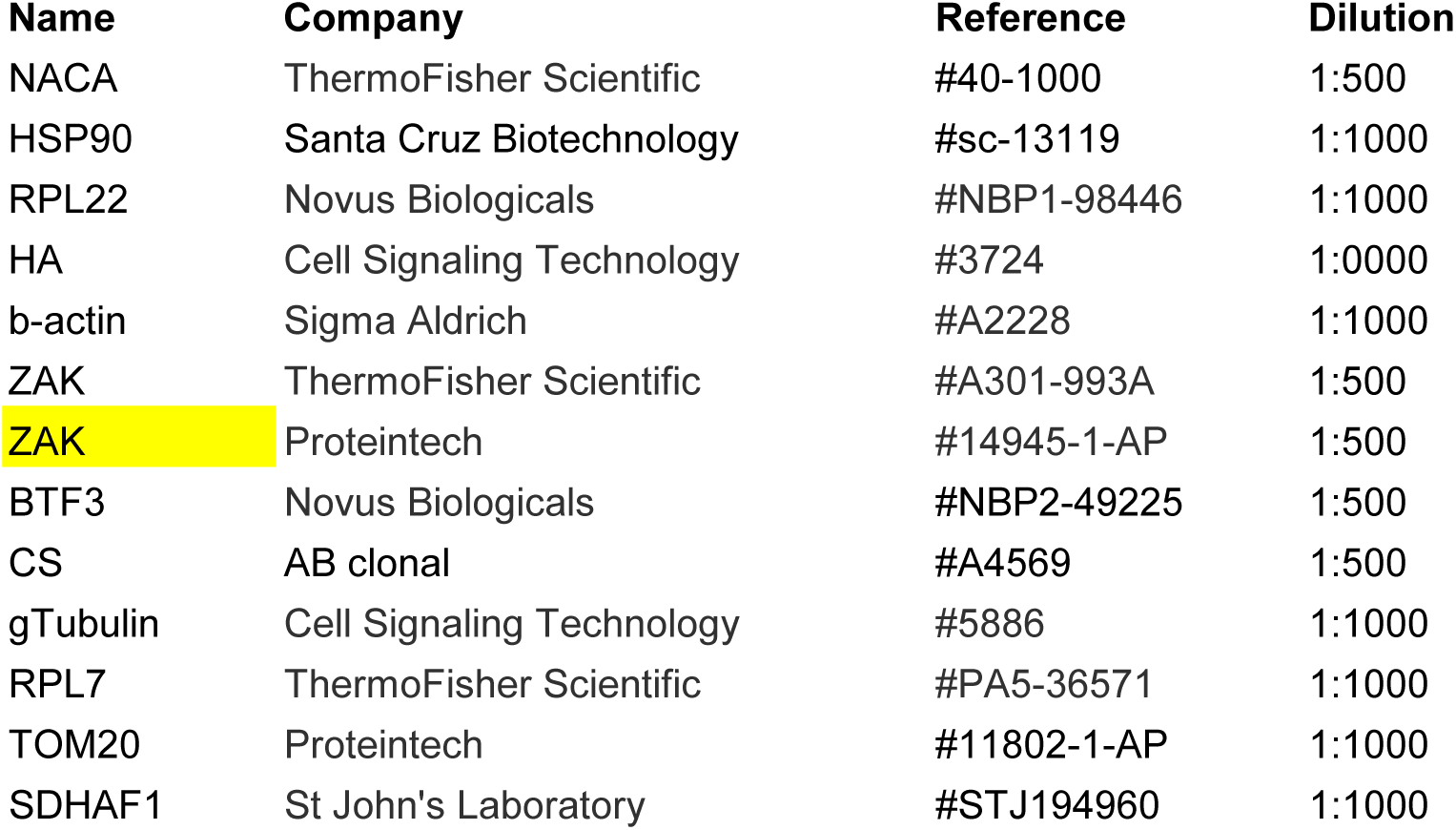

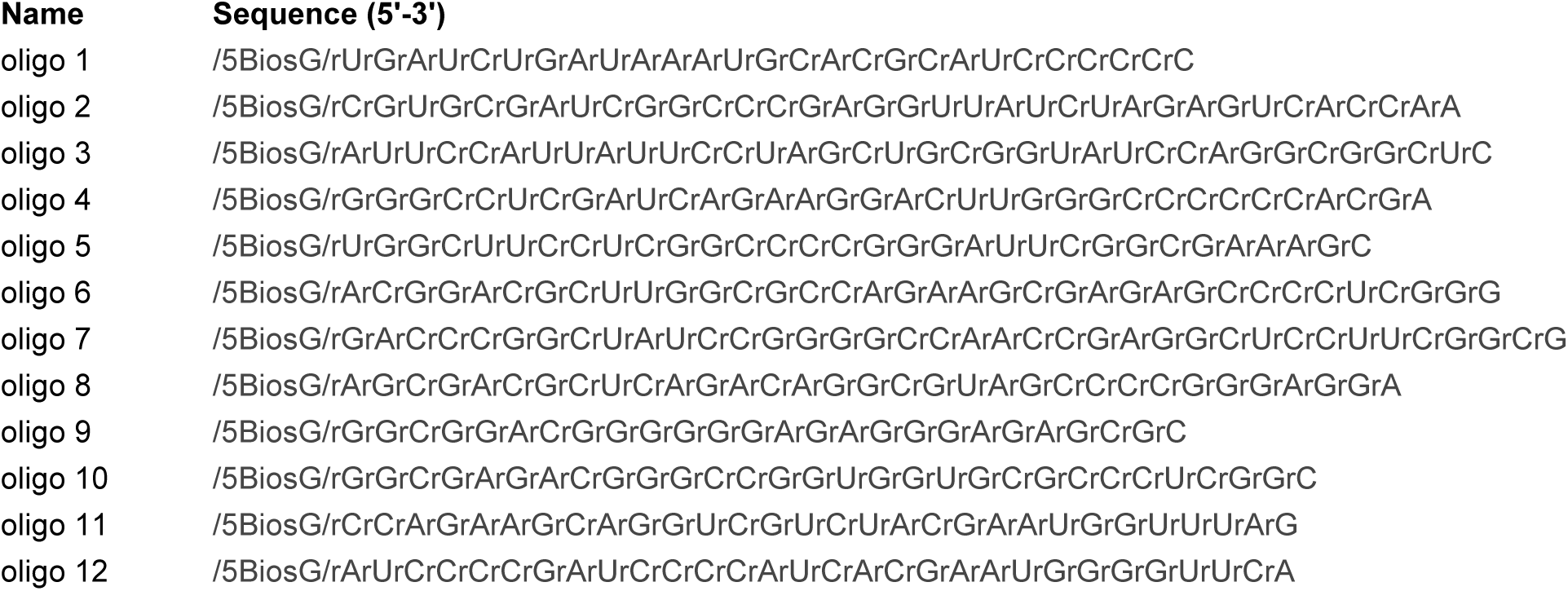

