## Supplementary Materials and Methods for "NAC-mediated ribosome localization regulates cell fate and metabolism in intestinal stem cells"

### Supp Fig 1

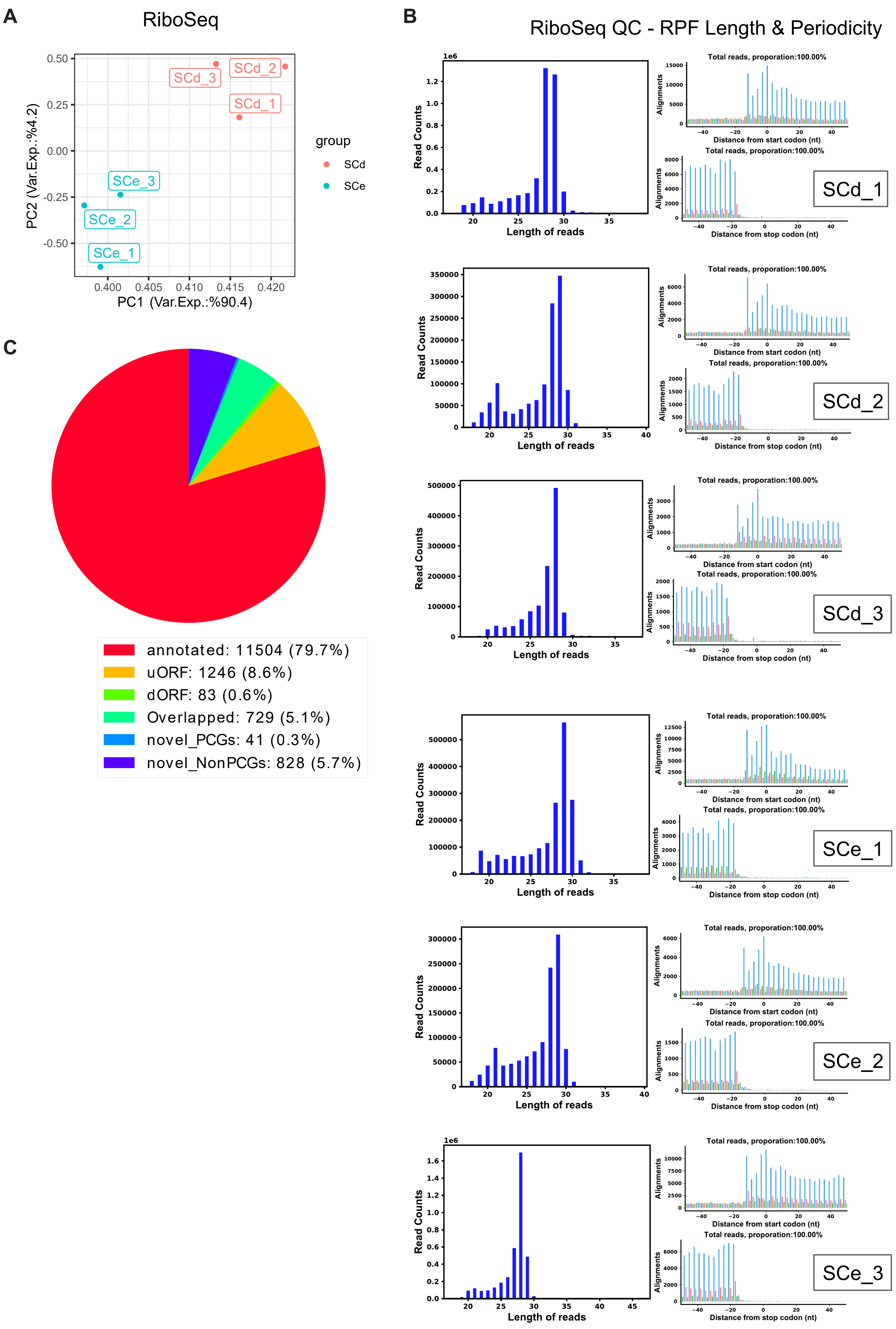
A
RiboSeq
PC1 (Var.Exp.:%90.4)
PC2 (Var.Exp.:%4.2)
group
SCd
SCe
SCd_1
SCd_2
SCd_3
SCe_1
SCe_2
SCe_3
B
RiboSeq QC - RPF Length & Periodicity
Read Counts
Length of reads
Total reads, proporation:100.00%
Alignments
Distance from start codon (nt)
Distance from stop codon (nt)
C
annotated: 11504 (79.7%)
uORF: 1246 (8.6%)
dORF: 83 (0.6%)
Overlapped: 729 (5.1%)
novel_PCGs: 41 (0.3%)
novel_NonPCGs: 828 (5.7%)

**Supplementary Figure 1 - Quality Control plots for Ribo-seq experiments performed in SCe and SCd organoids - Related to Figure 1**

A) PCA plot for RiboSeq performed in SCe and SCd organoids. Three biological replicates were used for each condition.

B) Quality control (QC) plots for RiboSeq experiments performed in SCe and SCd organoids, generated by the RiboCode tool. QC plots include read length histograms and periodicity plots of mRNA-mapped reads separately for each sample.

C) Pie chart depicting the RiboCode ORF prediction statistics using the RiboSeq data from all samples. Raw numbers and percentages of different classes of ORFs are presented.

### Supp Fig 2

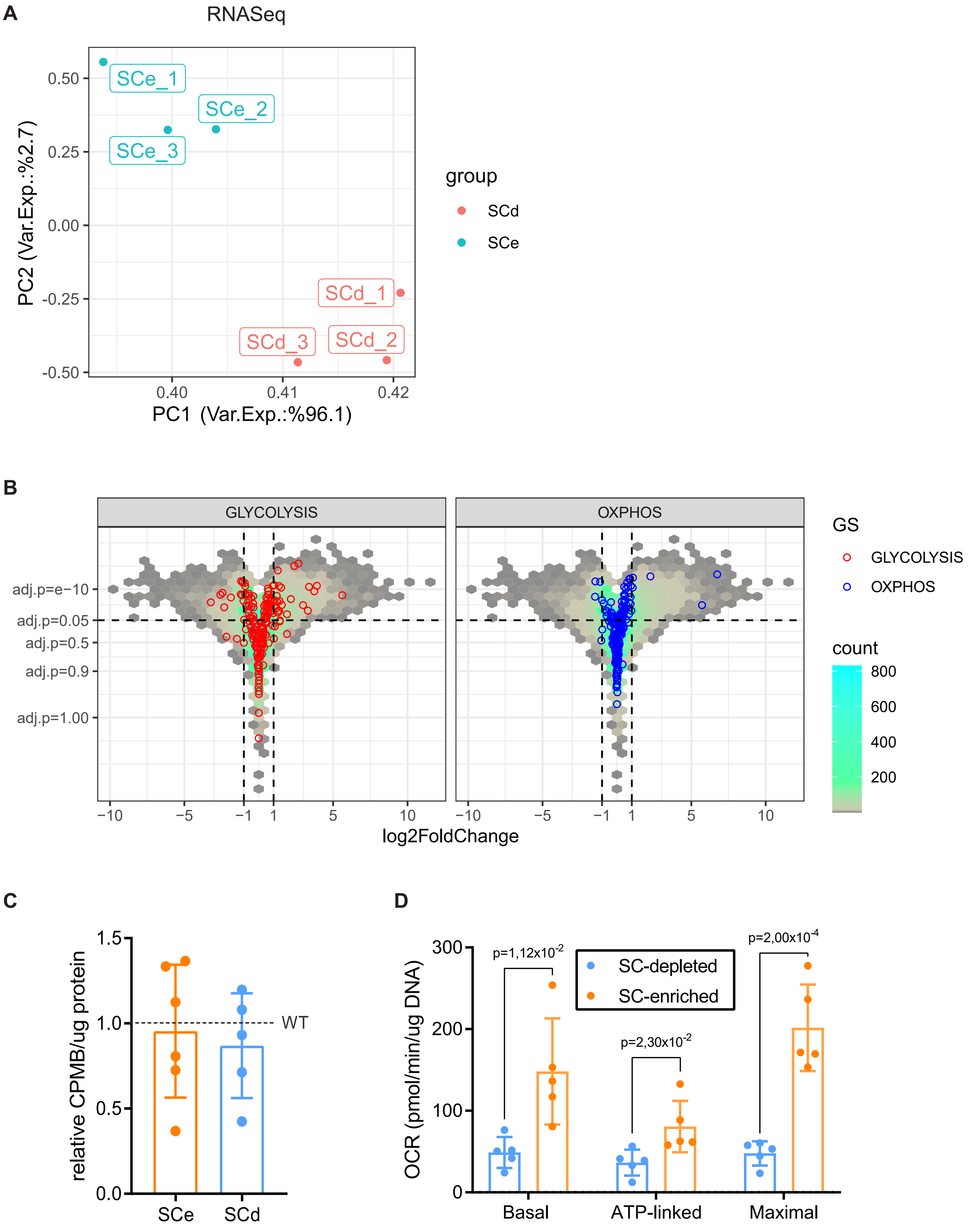
A
RNASeq
SCe_1
SCe_2
SCe_3
SCd_1
SCd_2
SCd_3
PC2 (Var.Exp.:%2.7)
PC1 (Var.Exp.:%96.1)
0.50
0.25
0.00
-0.25
-0.50
0.40
0.41
0.42
group
SCd
SCe
B
GLYCOLYSIS
OXPHOS
adj.p=e−10
adj.p=0.05
adj.p=0.5
adj.p=0.9
adj.p=1.00
−10
−5
−1
1
5
10
log2FoldChange
GS
GLYCOLYSIS
OXPHOS
count
800
600
400
200
C
relative CPMB/ug protein
1.5
1.0
0.5
0.0
WT
SCe
SCd
D
OCR (pmol/min/ug DNA)
300
200
100
0
p=1,12x10-2
p=2,30x10-2
p=2,00x10-4
SC-depleted
SC-enriched
Basal
ATP-linked
Maximal

**Supplementary Figure 2 - RNASeq analysis, total protein synthesis and seahorse quantification of SCe and SCd organoids - Related to Figure 1**

A) PCA plot for RNASeq performed in SCe and SCd organoids. Three biological replicates were used for each condition.

B) Volcano plot showing the differential expression results of SCe and SCd comparison, highlighting that most genes involved in glycolysis and OXPHOS do not seem to change significantly between the two conditions. p values were determined using the *DESEQ2* package.

C) Incorporation of $^{35}$S-methionine shows no differences in total protein synthesis between SCe and SCd cultures. Mean and SD are shown (n = 6 ( two biological replicates, each assessed in technical duplicates)). *p*-values were determined using a two-tailed *t*-test.

D) OCR analysis shows increased respiration in SCe compared to SCd organoids. Mean and SD are shown (n = 5 biological replicates). *p*-values were determined using a two-tailed t-test. Related to figure 1G.

#### Supp Fig 3

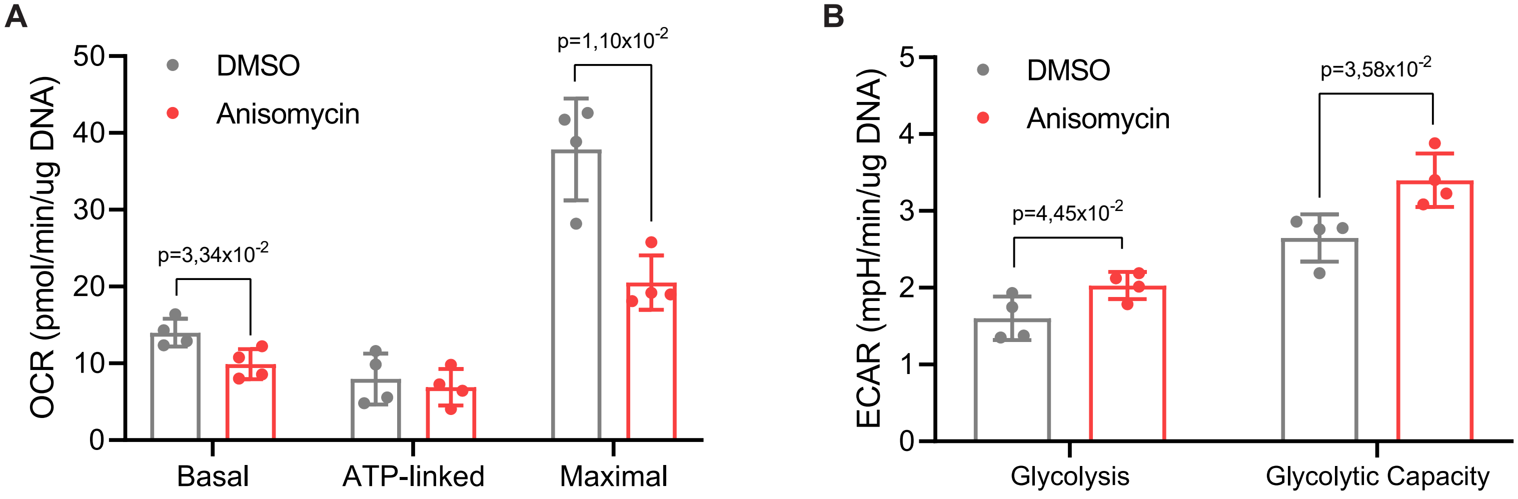
A
OCR (pmol/min/ug DNA)
0
10
20
30
40
50
DMSO
Anisomycin
p=3,34x10-2
p=1,10x10-2
Basal
ATP-linked
Maximal
B
ECAR (mpH/min/ug DNA)
0
1
2
3
4
5
DMSO
Anisomycin
p=4,45x10-2
p=3,58x10-2
Glycolysis
Glycolytic Capacity

**Supplementary Figure 3 - Seahorse analysis of WT organoids treated with anisomycin - Related to Figure 2**

A) OCR analysis shows decreased respiration in WT organoids following anisomycin treatment (1µM, 30 minutes). Mean and SD are shown (n = 4 biological replicates). *p*-values were determined using a two-tailed *t*-test. Related to Figure 2F.

B) ECAR analysis shows increased glycolytic rates after treating WT organoids with anisomycin (1µM, 30 minutes). Mean and SD are shown (n = 4 biological replicates). *p*-values were determined using a two-tailed *t*-test. Related to Figure 2G.

### Supp Fig 4

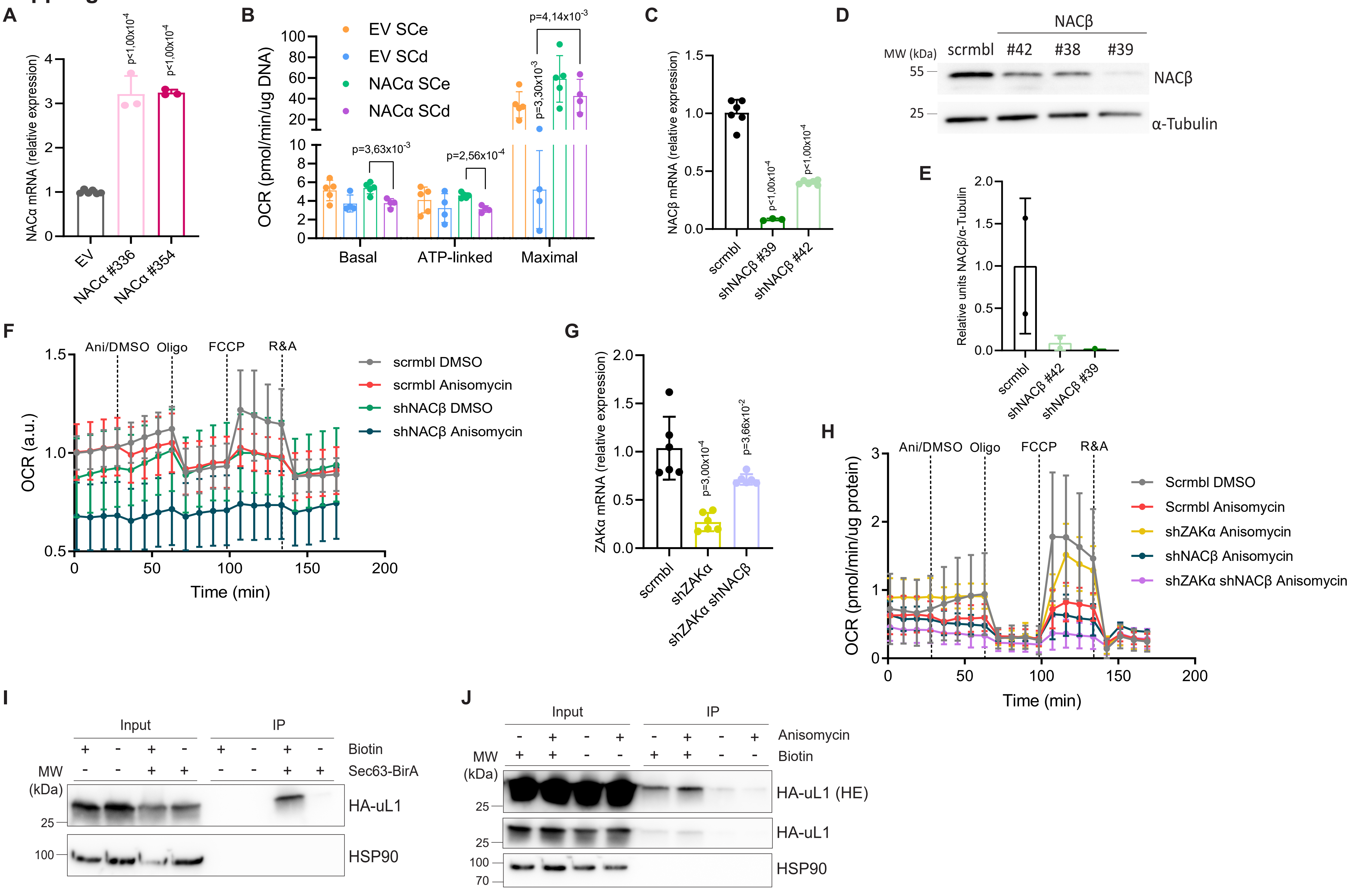
A
NACα mRNA (relative expression)
p<1,00x10-4
p<1,00x10-4
EV
NACα #336
NACα #354
B
OCR (pmol/min/ug DNA)
EV SCe
EV SCd
NACα SCe
NACα SCd
p=4,14x10-3
p=3,30x10-3
p=3,63x10-3
p=2,56x10-4
Basal
ATP-linked
Maximal
C
NACβ mRNA (relative expression)
p<1,00x10-4
p<1,00x10-4
scrmbl
shNACβ #39
shNACβ #42
D
NACβ
MW (kDa)
scrmbl
#42
#38
#39
55
25
NACβ
α-Tubulin
E
Relative units NACβ/α-Tubulin
scrmbl
shNACβ #42
shNACβ #39
F
Ani/DMSO
Oligo
FCCP
R&A
OCR (a.u.)
Time (min)
scrmbl DMSO
scrmbl Anisomycin
shNACβ DMSO
shNACβ Anisomycin
G
ZAKα mRNA (relative expression)
p=3,00x10-4
p=3,66x10-2
scrmbl
shZAKα
shZAKα shNACβ
H
Ani/DMSO
Oligo
FCCP
R&A
OCR (pmol/min/ug protein)
Time (min)
Scrmbl DMSO
Scrmbl Anisomycin
shZAKα Anisomycin
shNACβ Anisomycin
shZAKα shNACβ Anisomycin
I
Input
IP
Biotin
Sec63-BirA
MW (kDa)
25
100
HA-uL1
HSP90
J
Input
IP
Anisomycin
Biotin
MW (kDa)
25
25
100
70
HA-uL1 (HE)
HA-uL1
HSP90

**Supplementary Figure 4 - NAC overexpression and knock down effects on respiration rates and ribosome localization to the ER analysis upon NAC inhibition - Related to Figures 3 and 4**

A) RT-qPCR analysis of NACα expression in mouse intestinal organoids derived from two mice (#336 and #354) upon overexpression or transduction with empty vector (EV). *Hprt* was used as a housekeeping reference. Mean and SD are shown (*n* = 3 (one biological replicate assessed in technical triplicates)). *p*-values were determined using a two-tailed *t*-test.

B) OCR analysis shows a rescue of respiration rates in SCd organoids overexpressing NACα. Mean and SD are shown (n = 4 biological replicates). *p*-values were determined using a two-tailed *t*-test. Related to Figure 3A.

C) RT-qPCR analysis of NACβ expression in HCT116 cells upon knock down. β-*actin* was used as a housekeeping reference. Mean and SD are shown (*n* = 3 (one biological replicate assessed in technical triplicates) for shNACβ #39 and n = 6 (two biological replicates assessed in technical triplicates) for shNACβ #42). *p*-values were determined using a two-tailed *t*-test.

D) Western blot analysis of the levels of NACβ in HCT116 cells upon knock down. α-Tubulin was used as a loading control. Experiments were done in two biological replicates for shNACβ #42 and one for shNACβ #39.

E) Quantification of western blots shows a decrease in NACβ levels upon knock down. Mean and SD are shown (*n* = 1 biological replicate for shNACβ #39 and n = 2 biological replicates for shNACβ #42 all assessed in technical triplicates). *p*-values were determined using a two-tailed *t*-test

F) OCR analysis shows decreased respiration rates in WT HCT116 cells treated with anisomycin (1µM, 30 minutes), upon NACβ knockdown and when combining both anisomycin treatment with NACβ knockdown. Mean and SD are shown (n = 5 biological replicates for each of the 2 independent shRNAs). Related to Figure 3C.

G) RT-qPCR analysis of ZAKα expression HCT116 cells upon knock down. β-*actin* was used as a housekeeping reference. Mean and SD are shown (*n* = 6 (two biological replicates assessed in technical triplicates)). *p*-values were determined using a two-tailed *t*-test.

H) OCR analysis shows decreased respiration rates in WT HCT116 cells treated with anisomycin (1μM, 30 minutes) and the rescue of this decrease upon ZAKα knockdown. When NACβ is knocked down, the rescue of the anisomycin effect observed with the loss of ZAKα is not possible. Mean and SD are shown (n = 4 biological replicates). Related to Figure 3D.

I) Western blot analysis confirms the efficiency of pulling down biotinylated ribosomes, measured by HA-uL1 levels upon treating cells with biotin. Experiments were carried out in one biological replicate.

J) Western blot analysis shows an increase in the number of biotinylated ribosomes, accessed by HA-uL1 levels, following anisomycin treatment. HSP90 serves as a loading control. Experiments were carried out in one biological replicate.

| Name | Sequence (5'-3') |
|---|---|
| mHprt_fw | CTGGTGAAAGGACCTCTCG |
| mHprt_rev | TGAAGTACTCATTATAGTCAAGGGCA |
| mLgr5_fw | ACCCGCCAGTCTCCTACATC |
| mLgr5_rev | GCATCTAGGCGCAGGGATTG |
| mAxin2_fw | GCGACGCACTGACCGACGAT |
| mAxin2_rev | GCAGGCGGTGGGTTCTCGGA |
| mLyz1_fw | CGGTTTTGACATTGTGTTCGC |
| mLyz1_rev | GAGACCGAAGCACCGACTATG |
| mAlpi_fw | ATGATGCCAACCGAAACCCC |
| mAlpi_rev | GCGTGTCCTTCTCATTGGTAA |
| mMex3a_fw | ACACCACGGAGTGCGTTC |
| mMex3a_rev | GTTGGTTTTGGCCCTCAGA |
| hACTB_fw | ACCTTCTACAATGAGCTGCG |
| hACTB_rev | CCTGGATAGCAACGTACATGG |
| hNACB_fw | CAGGAGCCTCAGATGAAAGAAA |
| hNACB_rev | TCTCAGCATGGCCTGTAATG |
| hZAKα_fw | CCACTCCTTCAAGAGGAAGATAC |
| hZAKα_rev | CCCTCTGGCTCTCTCTTTATTG |
| hZAK_fw | CAAGGAGGTGGCTGTAAAGAA |
| hZAK_rev | ACACAGGGAGACTCTGGATAA |
| mNACA NM013608.3 _fw | GAAGGCAAGGAAGGCTATGT |
| mNACA NM013608.3_rev | CGGTTGGAGTCTGAGTGTTT |
| hMdh2_fw | TGAAGAACAGCCCCTTGGTG |
| hMdh2_rev | GGTCCGAGGTAGCCTTTCAC |
| hPiC_fw | AGGATGGTGTTCGTGGTTTG |
| hPiC_rev | TGTGCGCCAGAGATAAGTATTC |
| hAnt1_fw | AGGGTTTCAACGTCTCTGTC |
| hAnt1_rev | GTCACACTCTGGGCAATCAT |
| hAtp5b_fw | TTGGTCCTGAGACTTTGGGC |
| hAtp5b_rev | CCTCAGCATGAATGGGAGCA |
| hCi30_fw | GATGAAGTGAAGCGGGTGGT |
| hCi30_rev | GGCGATAGACTGGGAAAGCC |
| hCox6c_fw | ATGGCTCCCGAAGTTTTGCC |
| hCox6c_rev | CCCCAGGGATAGCACGAATG |
| hAdh5_fw | GGCTCATGAAGTTCGAATCAAG |
| hAdh5_rev | ACTCCCTCACCAACACTTTC |
| hAtp5a1_fw | GATCCGCTGCCCAAACC |
| hAtp5a1_rev | GCCAATTCCAGCTTCATGGT |
| hSdhaf1_fw | GCCTTTTCCGATGCTTGGAG |
| hSdhaf1_rev | CCGTACCTTCGGGCAAGTAG |
| h18S rRNA_fw | GTAACCCGTTGAACCCCATT |
| h18S rRNA_rev | CCATCCAATCGGTAGTAGGC |
| RTP_RiboSeq | GCCTTGGCACCCGAGAATTCCA |
| RP1_RiboSeq | AATGATACGGCGACCACCGAGATCTACACGTTCAGAGTTCTACAGTCCGA |
| RPI1_RiboSeq | CAAGCAGAAGACGGCATACGAGATCGTGATGTGACTGGAGTTCCTTGGCACCCGAGAAT |
| RPI2_RiboSeq | CAAGCAGAAGACGGCATACGAGATACATCGGTGACTGGAGTTCCTTGGCACCCGAGAAT |
| RPI3_RiboSeq | CAAGCAGAAGACGGCATACGAGATGCCTAAGTGACTGGAGTTCCTTGGCACCCGAGAAT |
| RPI4_RiboSeq | CAAGCAGAAGACGGCATACGAGATTGGTCAGTGACTGGAGTTCCTTGGCACCCGAGAAT |
| RPI5_RiboSeq | CAAGCAGAAGACGGCATACGAGATCACTGTGTGACTGGAGTTCCTTGGCACCCGAGAAT |
| RPI6_RiboSeq | CAAGCAGAAGACGGCATACGAGATATTGGCGTGACTGGAGTTCCTTGGCACCCGAGAAT |
| PC1_fw_CRISPR screen | TAAAGGGACGGTTAATCGCTCACC |
| PC1_rev_CRISPR screen | ACACTCCAATGGGAGACGTTGTC |
| PCR1_fw_BC01 | ACACTCTTTCCCTACACGACGCTCTTCCGATCTCGTGATGGCTTTATATATCTTGTGGAAA |
| PCR1_fw_BC02 | ACACTCTTTCCCTACACGACGCTCTTCCGATCTACATCGGGCTTTATATATCTTGTGGAAA |
| PCR1_fw_BC03 | ACACTCTTTCCCTACACGACGCTCTTCCGATCTGCCTAAGGCTTTATATATCTTGTGGAAA |
| PCR1_fw_BC04 | ACACTCTTTCCCTACACGACGCTCTTCCGATCTTGGTCAGGCTTTATATATCTTGTGGAAA |
| PCR1_fw_BC05 | ACACTCTTTCCCTACACGACGCTCTTCCGATCTCACTGTGGCTTTATATATCTTGTGGAAA |
| PCR1_fw_BC06 | ACACTCTTTCCCTACACGACGCTCTTCCGATCTATTGGCGGCTTTATATATCTTGTGGAAA |
| PCR1_rev_lentiguide | GTGACTGGAGTTCAGACGTGTGCTCTTCCGATCTACTGACGGGCACCGGAGCCAATTCC |
| PCR2_fw_P5 | AATGATACGGCGACCACCGAGATCTACACTCTTTCCCTACACGACGCTCTTCCGATCT |
| PCR2_rev_P7_BC01 | CAAGCAGAAGACGGCATACGAGATATCACGGTGACTGGAGTTCAGACGTGTGCTCTTCC |
| PCR2_rev_P7_BC02 | CAAGCAGAAGACGGCATACGAGATCGATGTGTGACTGGAGTTCAGACGTGTGCTCTTCC |
| PCR2_rev_P7_BC03 | CAAGCAGAAGACGGCATACGAGATTTAGGCGTGACTGGAGTTCAGACGTGTGCTCTTCC |
| PCR2_rev_P7_BC04 | CAAGCAGAAGACGGCATACGAGATTGACCAGTGACTGGAGTTCAGACGTGTGCTCTTCC |
| PCR2_rev_P7_BC05 | CAAGCAGAAGACGGCATACGAGATACATGTGTGACTGGAGTTCAGACGTGTGCTCTTCC |
| PCR2_rev_P7_BC06 | CAAGCAGAAGACGGCATACGAGATGCCAATGTGACTGGAGTTCAGACGTGTGCTCTTCC |

| Name | Company | Reference | Dilution |
| --- | --- | --- | --- |
| NACA | ThermoFisher Scientific | #40-1000 | 1:500 |
| HSP90 | Santa Cruz Biotechnology | #sc-13119 | 1:1000 |
| RPL22 | Novus Biologicals | #NBP1-98446 | 1:1000 |
| HA | Cell Signaling Technology | #3724 | 1:0000 |
| b-actin | Sigma Aldrich | #A2228 | 1:1000 |
| ZAK | ThermoFisher Scientific | #A301-993A | 1:500 |
| ZAK | Proteintech | #14945-1-AP | 1:500 |
| BTF3 | Novus Biologicals | #NBP2-49225 | 1:500 |
| CS | AB clonal | #A4569 | 1:500 |
| gTubulin | Cell Signaling Technology | #5886 | 1:1000 |
| RPL7 | ThermoFisher Scientific | #PA5-36571 | 1:1000 |
| TOM20 | Proteintech | #11802-1-AP | 1:1000 |
| SDHAF1 | St John's Laboratory | #STJ194960 | 1:1000 |

| Name | Sequence (5'-3') |
| --- | --- |
| oligo 1 | /5BiosG/rUrGrArUrCrUrGrArUrArArArUrGrCrArCrGrCrArUrCrCrCrCrC |
| oligo 2 | /5BiosG/rCrGrUrGrCrGrArUrCrGrGrCrCrCrGrArGrGrUrUrArUrCrUrArGrArGrUrCrArCrCrArA |
| oligo 3 | /5BiosG/rArUrUrCrCrArUrUrArUrUrCrCrUrArGrCrUrGrCrGrGrUrArUrCrCrArGrGrCrGrGrCrUrC |
| oligo 4 | /5BiosG/rGrGrGrCrCrUrCrGrArUrCrArGrArArGrGrArCrUrUrGrGrGrCrCrCrCrCrCrArCrGrA |
| oligo 5 | /5BiosG/rUrGrGrCrUrUrCrCrUrCrGrGrCrCrCrCrGrGrGrArUrUrCrGrGrCrGrArArArGrC |
| oligo 6 | /5BiosG/rArCrGrGrArCrGrCrUrUrGrGrCrGrCrCrArGrArArGrCrGrArGrArGrCrCrCrCrUrCrGrGrG |
| oligo 7 | /5BiosG/rGrArCrCrCrGrGrCrUrArUrCrCrGrGrGrGrCrCrArArCrCrGrArGrGrCrUrCrCrUrUrCrGrGrCrG |
| oligo 8 | /5BiosG/rArGrCrGrArCrGrCrUrCrArGrArCrArGrGrCrGrUrArGrCrCrCrCrGrGrGrArGrGrA |
| oligo 9 | /5BiosG/rGrGrCrGrGrArCrGrGrGrGrGrArGrArGrGrGrArGrArGrCrGrC |
| oligo 10 | /5BiosG/rGrGrCrGrArGrArCrGrGrGrCrCrGrGrUrGrGrUrGrCrGrCrCrCrUrCrGrGrC |
| oligo 11 | /5BiosG/rCrCrArGrArArGrCrArGrGrUrCrGrUrCrUrArCrGrArArUrGrGrUrUrUrArG |
| oligo 12 | /5BiosG/rArUrCrCrCrCrGrArUrCrCrCrCrArUrCrArCrGrArArUrGrGrGrGrUrUrCrA |
