## Supplementary Figure 1 for "NAC-mediated ribosome localization regulates cell fate and metabolism in intestinal stem cells"

**Supp Fig 1**

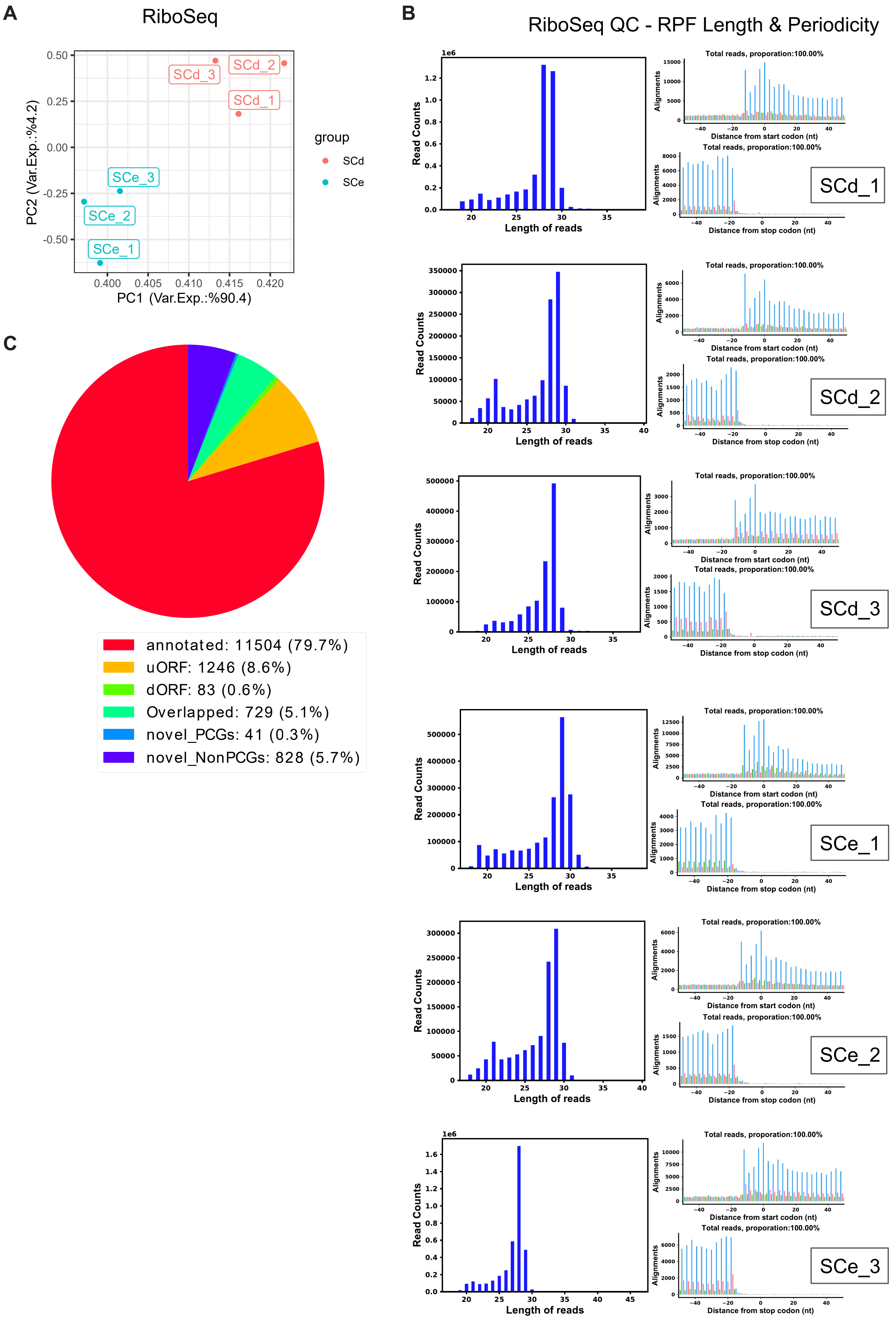

A
RiboSeq
PC1 (Var.Exp.:%90.4)
PC2 (Var.Exp.:%4.2)
group
SCd
SCe
SCd_1
SCd_2
SCd_3
SCe_1
SCe_2
SCe_3
B
RiboSeq QC - RPF Length & Periodicity
Read Counts
Length of reads
Total reads, proporation:100.00%
Alignments
Distance from start codon (nt)
Distance from stop codon (nt)
C
annotated: 11504 (79.7%)
uORF: 1246 (8.6%)
dORF: 83 (0.6%)
Overlapped: 729 (5.1%)
novel_PCGs: 41 (0.3%)
novel_NonPCGs: 828 (5.7%)

**Supplementary Figure 1 - Quality Control plots for Ribo-seq experiments performed in SCe and SCd organoids - Related to Figure 1**

A) PCA plot for RiboSeq performed in SCe and SCd organoids. Three biological replicates were used for each condition.

B) Quality control (QC) plots for RiboSeq experiments performed in SCe and SCd organoids, generated by the RiboCode tool. QC plots include read length histograms and periodicity plots of mRNA-mapped reads separately for each sample.

C) Pie chart depicting the RiboCode ORF prediction statistics using the RiboSeq data from all samples. Raw numbers and percentages of different classes of ORFs are presented.
