## Supplementary Figure 2 for "NAC-mediated ribosome localization regulates cell fate and metabolism in intestinal stem cells"

Supp Fig 2

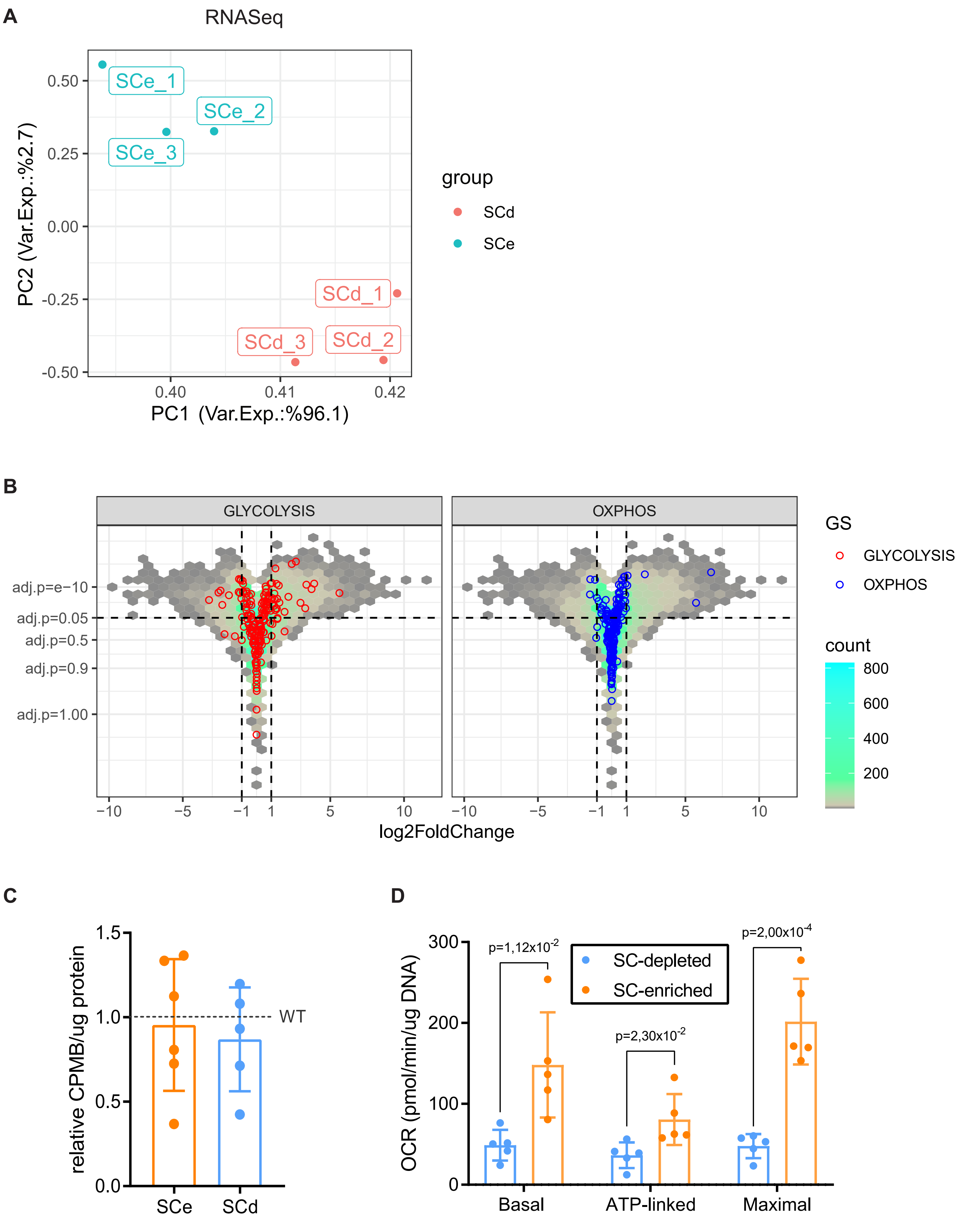
A
RNASeq
SCe_1
SCe_2
SCe_3
SCd_1
SCd_2
SCd_3
0.50
0.25
0.00
-0.25
-0.50
0.40
0.41
0.42
PC2 (Var.Exp.:%2.7)
PC1 (Var.Exp.:%96.1)
group
SCd
SCe
B
GLYCOLYSIS
OXPHOS
adj.p=e−10
adj.p=0.05
adj.p=0.5
adj.p=0.9
adj.p=1.00
−10
−5
−1
1
5
10
log2FoldChange
GS
GLYCOLYSIS
OXPHOS
count
800
600
400
200
C
1.5
1.0
0.5
0.0
WT
relative CPMB/ug protein
SCe
SCd
D
300
200
100
0
OCR (pmol/min/ug DNA)
p=1,12x10-2
p=2,30x10-2
p=2,00x10-4
SC-depleted
SC-enriched
Basal
ATP-linked
Maximal

**Supplementary Figure 2 - RNASeq analysis, total protein synthesis and seahorse quantification of SCe and SCd organoids - Related to Figure 1**

A) PCA plot for RNASeq performed in SCe and SCd organoids. Three biological replicates were used for each condition.

B) Volcano plot showing the differential expression results of SCe and SCd comparison, highlighting that most genes involved in glycolysis and OXPHOS do not seem to change significantly between the two conditions. p values were determined using the *DESEQ2* package.

C) Incorporation of $^{35}$S-methionine shows no differences in total protein synthesis between SCe and SCd cultures. Mean and SD are shown (n = 6 ( two biological replicates, each assessed in technical duplicates)). *p*-values were determined using a two-tailed *t*-test.

D) OCR analysis shows increased respiration in SCe compared to SCd organoids. Mean and SD are shown (n = 5 biological replicates). *p*-values were determined using a two-tailed t-test. Related to figure 1G.
