## Supplementary Figure 3 for "NAC-mediated ribosome localization regulates cell fate and metabolism in intestinal stem cells"

### Supp Fig 3

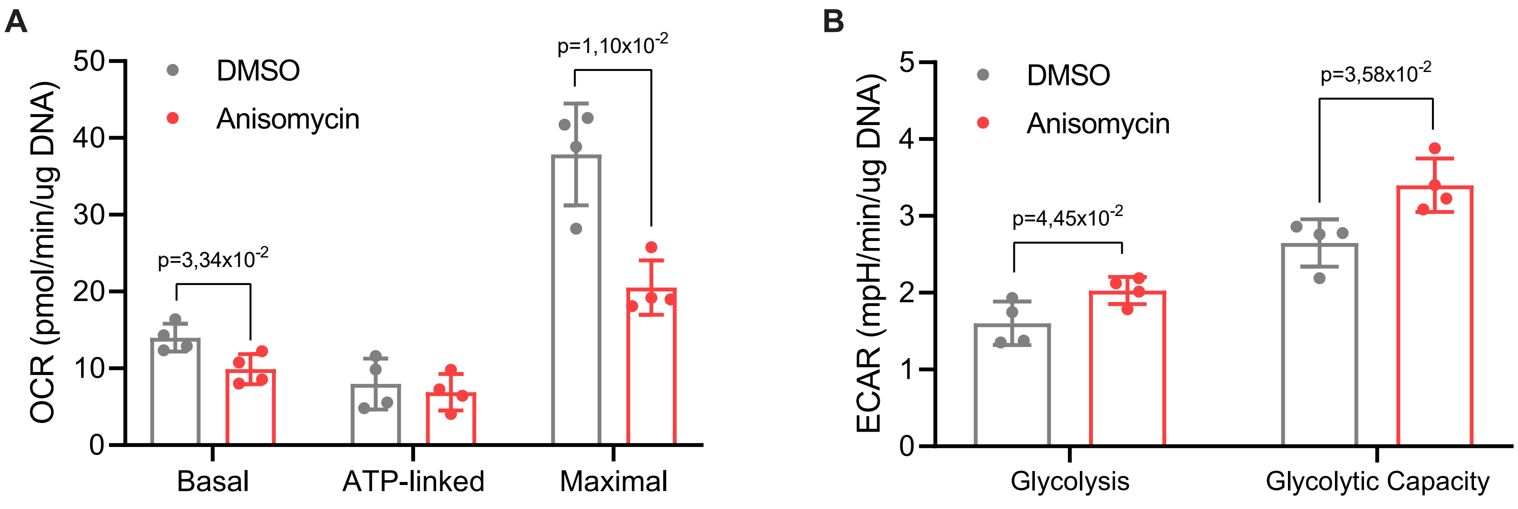
A
OCR (pmol/min/ug DNA)
0
10
20
30
40
50
DMSO
Anisomycin
p=3,34x10-2
p=1,10x10-2
Basal
ATP-linked
Maximal
B
ECAR (mpH/min/ug DNA)
0
1
2
3
4
5
DMSO
Anisomycin
p=4,45x10-2
p=3,58x10-2
Glycolysis
Glycolytic Capacity

**Supplementary Figure 3 - Seahorse analysis of WT organoids treated with anisomycin - Related to Figure 2**

A) OCR analysis shows decreased respiration in WT organoids following anisomycin treatment (1μM, 30 minutes). Mean and SD are shown (n = 4 biological replicates). *p*-values were determined using a two-tailed *t*-test. Related to Figure 2F.

B) ECAR analysis shows increased glycolytic rates after treating WT organoids with anisomycin (1μM, 30 minutes). Mean and SD are shown (n = 4 biological replicates). *p*-values were determined using a two-tailed *t*-test. Related to Figure 2G.
