## Supplementary Figure 4 for "NAC-mediated ribosome localization regulates cell fate and metabolism in intestinal stem cells"

Supp Fig 4

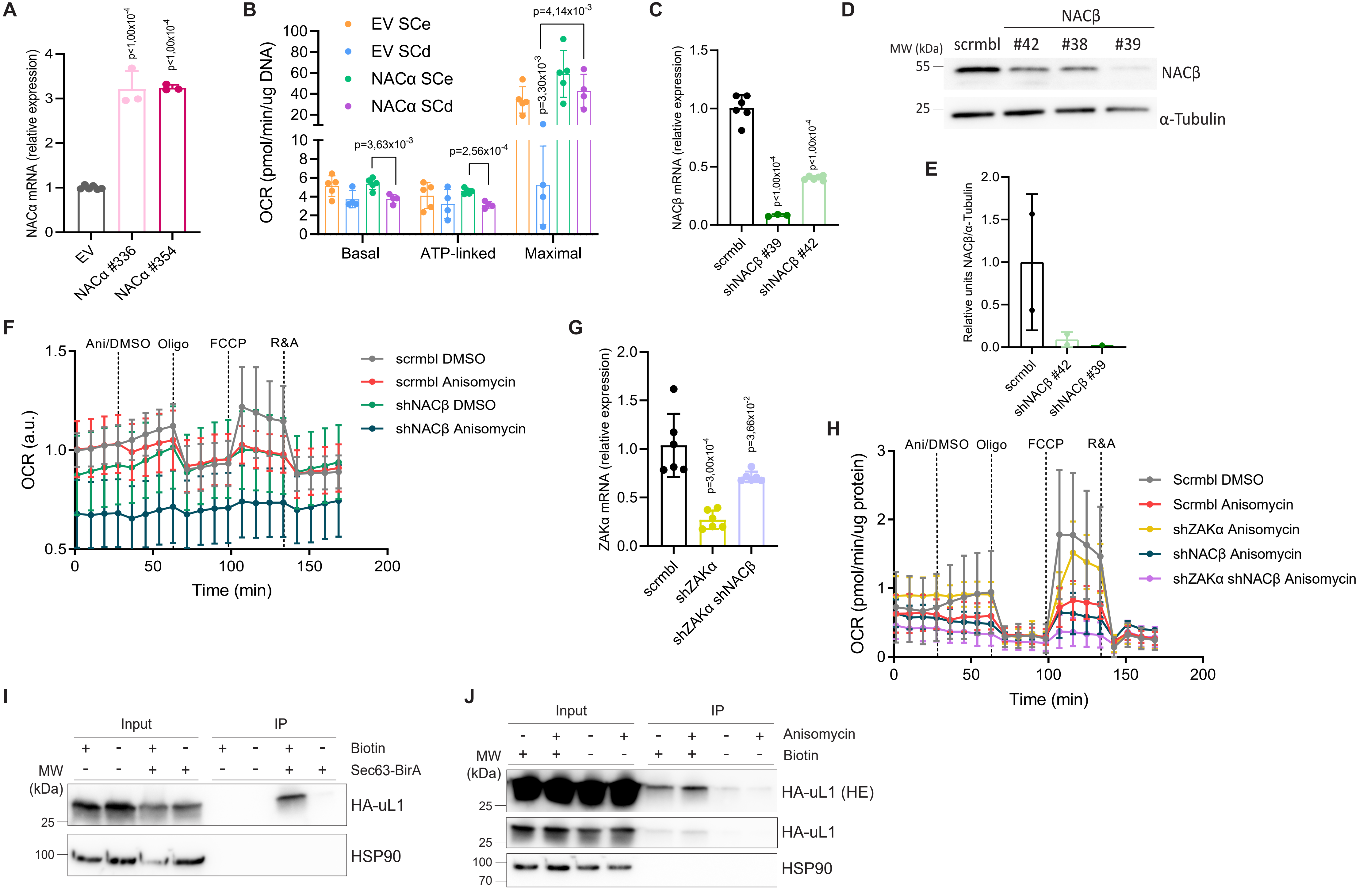

A
NACα mRNA (relative expression)
p<1,00x10^-4
p<1,00x10^-4
EV
NACα #336
NACα #354
B
OCR (pmol/min/ug DNA)
EV SCe
EV SCd
NACα SCe
NACα SCd
p=4,14x10^-3
p=3,30x10^-3
p=3,63x10^-3
p=2,56x10^-4
Basal
ATP-linked
Maximal
C
NACβ mRNA (relative expression)
p<1,00x10^-4
p<1,00x10^-4
scrmbl
shNACβ #39
shNACβ #42
D
NACβ
MW (kDa)
scrmbl
#42
#38
#39
55
25
NACβ
α-Tubulin
E
Relative units NACβ/α-Tubulin
scrmbl
shNACβ #42
shNACβ #39
F
Ani/DMSO
Oligo
FCCP
R&A
OCR (a.u.)
Time (min)
scrmbl DMSO
scrmbl Anisomycin
shNACβ DMSO
shNACβ Anisomycin
G
ZAKα mRNA (relative expression)
p=3,00x10^-4
p=3,66x10^-2
scrmbl
shZAKα
shZAKα shNACβ
H
Ani/DMSO
Oligo
FCCP
R&A
OCR (pmol/min/ug protein)
Time (min)
Scrmbl DMSO
Scrmbl Anisomycin
shZAKα Anisomycin
shNACβ Anisomycin
shZAKα shNACβ Anisomycin
I
Input
IP
Biotin
Sec63-BirA
MW (kDa)
25
100
HA-uL1
HSP90
J
Input
IP
Anisomycin
Biotin
MW (kDa)
25
25
100
70
HA-uL1 (HE)
HA-uL1
HSP90
